# Piggybacking functionalized DNA nanostructures into live cell nuclei

**DOI:** 10.1101/2023.12.30.573746

**Authors:** Golbarg M. Roozbahani, Patricia Colosi, Attila Oravecz, Elena M. Sorokina, Wolfgang Pfeifer, Siamak Shokri, Yin Wei, Pascal Didier, Marcello DeLuca, Gaurav Arya, László Tora, Melike Lakadamyali, Michael G. Poirier, Carlos E. Castro

## Abstract

DNA origami (DO) are promising tools for *in vitro* or *in vivo* applications including drug delivery; biosensing, detecting biomolecules; and probing chromatin sub-structures. Targeting these nanodevices to mammalian cell nuclei could provide impactful approaches for probing visualizing and controlling important biological processes in live cells. Here we present an approach to deliver DO strucures into live cell nuclei. We show that labelled DOs do not undergo detectable structural degradation in cell culture media or human cell extracts for 24 hr. To deliver DO platforms into the nuclei of human U2OS cells, we conjugated 30 nm long DO nanorods with an antibody raised against the largest subunit of RNA Polymerase II (Pol II), a key enzyme involved in gene transcription. We find that DOs remain structurally intact in cells for 24hr, including within the nucleus. Using fluorescence microscopy we demonstrate that the electroporated anti-Pol II antibody conjugated DOs are efficiently piggybacked into nuclei and exihibit sub-diffusive motion inside the nucleus. Our results reveal that functionalizing DOs with an antibody raised against a nuclear factor is a highly effective method for the delivery of nanodevices into live cell nuclei.

## INTRODUCTION

Recent advances in DNA nanotechnology have presented promising opportunities for applications in areas like drug delivery, biosensing, and biomanufacturing [1]–[3]. In particular, DNA origami (DO),[4] where a long template strand is folded into a compact shape by base-pairing with many shorter strands, enables fabrication of nanostructures with complex and precise shape, custom functionalization, and tunable mechanical properties[5], [6]. These features make DO devices attractive as platforms for targeted therapies,[7] biophysical measurements[8], or controlling molecular interactions[9], [10]. Many of these applications either require or can be enhanced by effective methods to deliver DO into intracellular environments. Prior studies have demonstrated uptake of DO into cells[11]–[13], but the trafficking of DOs upon entry into live cells and specifically to nuclei is less well-understood and/or developed. Methods for the efficient delivery of DOs into live cell nuclei could greatly enhance existing applications in therapeutic delivery, for example gene delivery,[14]–[16] and could enable translation of other functions of DO like biophysical measurement or imaging into cell nuclei.

The nucleus houses the cell’s genetic material and the machinery essential for transcription and other processes vital to gene expression and regulation[17], [18]. Consequently, targeting molecular structures and devices to the nucleus is an attractive approach for many therapies and may present opportunities for nanoscale tools to probe or control the genetic or epigenetic processes that regulate cell function. For example, recent *in vitro* work has demonstrated nanodevices as tools for sequestering or organizing biomolecules or larger complexes,[19]–[21] imaging biomolecules at high resolution,[22], [23] and manipulating enzymatic reactions,[24], [25] all of which could be useful inside cells and cellular compartments. Delivering DO nanodevices to cell nuclei is attractive for applications like nucleic acid detection[26], [27], biophysical probing of chromatin sub-structures (previously demonstrated *in vitro* [28], [29]), and gene delivery.[14]–[16]

While significant efforts have studied the delivery and uptake of DO nanostructures into live cells,[11], [13], [30], [31] only recently has the specific delivery of DO structures to the nucleus been explored, focused in the context of gene delivery.[14]–[16], [32] These studies have established DO as useful tool for the delivery of genetic information into live cells. Even though these prior studies focused on gene expression, key questions remain unclear: i) are these DO structures stable inside the cell?, ii) how many of the DO structures reach the nuclei, and iii) can intact DOs can be delivered into the nucleus? Hence, there remains a critical need for robust methods to deliver DO nanostructures to live cell nuclei, which would be an essential step to enabling intranuclear functions that rely on the structure and not just the encoded sequence.

Here, we present a novel approach for the delivery of intact DO nanstructures into live cells and specifically to the nucleus (Figure 1). Inspired by recent work focused on the delivery of antibodies into live cell nuclei,[33]–[35] our method involves the conjugation of DO nanostructures to antibodies that bind to neosynthetized proteins in the cytoplasm, which function in the nucleus and thus naturally cycle to the nucleus, thereby carrying, or "piggybacking," the DOs along with them. We chose the large subunit of RNA polymerase Pol II, a pivotal enzyme involved in gene transcription, as a molecule to target the neosynthetized subunit in the cytoplasm. Our prior work demonstrated that the piggybacking approach is effective for the delivery of antibodies with high affinity towards Pol II into live cell nuclei.[33] Here we show that, after electroporation into the ctyoplasm, Pol II antibody-conjugated 30 nm nanorod DO structures can enter the nuclei of U2OS cells, as confirmed by fluorescence microscopy, and exhibit sub-diffusive motion within live cell nuclei. We also studied the stability of DO in cell culture media and different cell lysates using gel electrophoresis and transmission electron microscopy (TEM), and inside live cells using fluorescence imaging. These analyses reveal the structural integrity of the DO over extended periods in cell media and extracts, and confirm that DOs remain structurally stable 24 hr after electroporation both in the cytoplasm and after piggybacking into the nucleus. Combined, our results establish a basis to implement DO nanodevices as tools for imaging, detection, biophysical measurements, or other applications inside cell nuclei.

**Figure 1:**
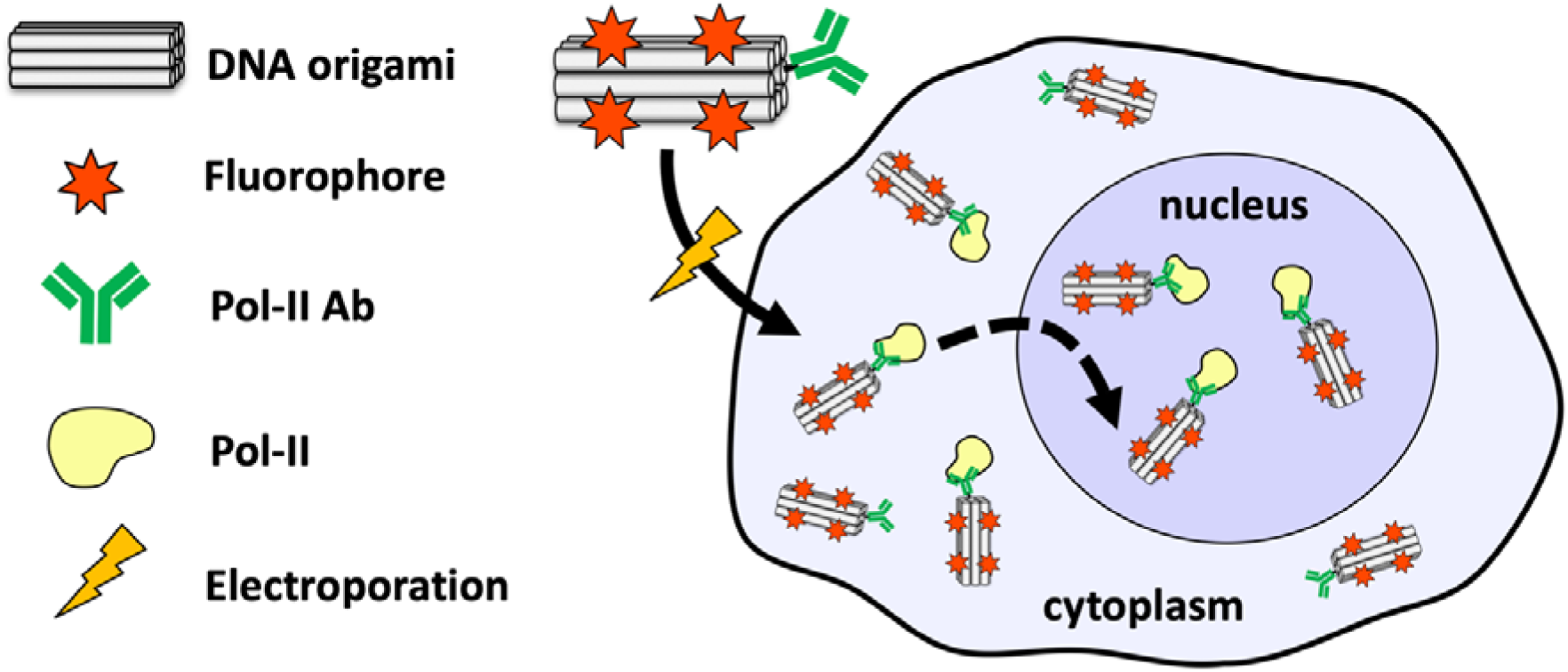
Concept for piggybacking DNA origami nanostructures into the nucleus. DNA origami nanostructures functionalized with RNA polymerase Pol II targeting antibodies and 8 Cy5 fluorophores are electroporated into cells, bind to Pol II, and then are imported or piggybacked into the nucleus.

## MATERIAL AND METHODS

### Design and simulation of DNA origami nanostructures

To achive the delivery of DOs to the nuclei of live cells, we initially tested two DO designs, an 8-helix bundle (8HB), which is 30 nm long with a molecular weight of ∼0.5 MDa, and a 26-helix bundle (26HB), which is approximately 90 nm long with a molecular weight of ∼5 MDa. Prior research has demonstrated the efficient folding and stability of the 26HB in cell culture media, as well as its effective cellular uptake,[36], [37] and the 8HB design uses a similar, but smaller square lattice cross-section. We used a previously reported design for the 26HB structures [36]–[38]. The 8HB nanostructure was designed in caDNAno[39] (Supplementary Figure S1 and Table S1, design available on nanobase.org), using a hollow square-lattice cross-section.[40] The staple strand routing was designed to contain ideally one long continuous duplex region per strand, which has been shown to facilitate robust folding.[41], [42] The scaffold routing was designed to contain a seam near the middle of the bundle, which has been shown to inhibit isomerization of the structure.[4], [43] Coarse grained MD simulations were performed using the oxDNA model,[44]–[46] after converting the caDNAno output files through tacoxDNA[47] into oxDNA topology and configuration files. Initial relaxation was performed using default parameters (oxdna.org). Simulations were run for 100,000,000 steps, and the mean over the full trajectory was used to depict the 8-helix bundle structure in figures 2A and 2C. For the depiction of 8HB with overhangs and antibody, the relaxed structure was converted to an all-atom PDB representation and visualized alongside a PDB representation of antibody (1igt) in ChimeraX[48] for scale.

**Figure 2:**
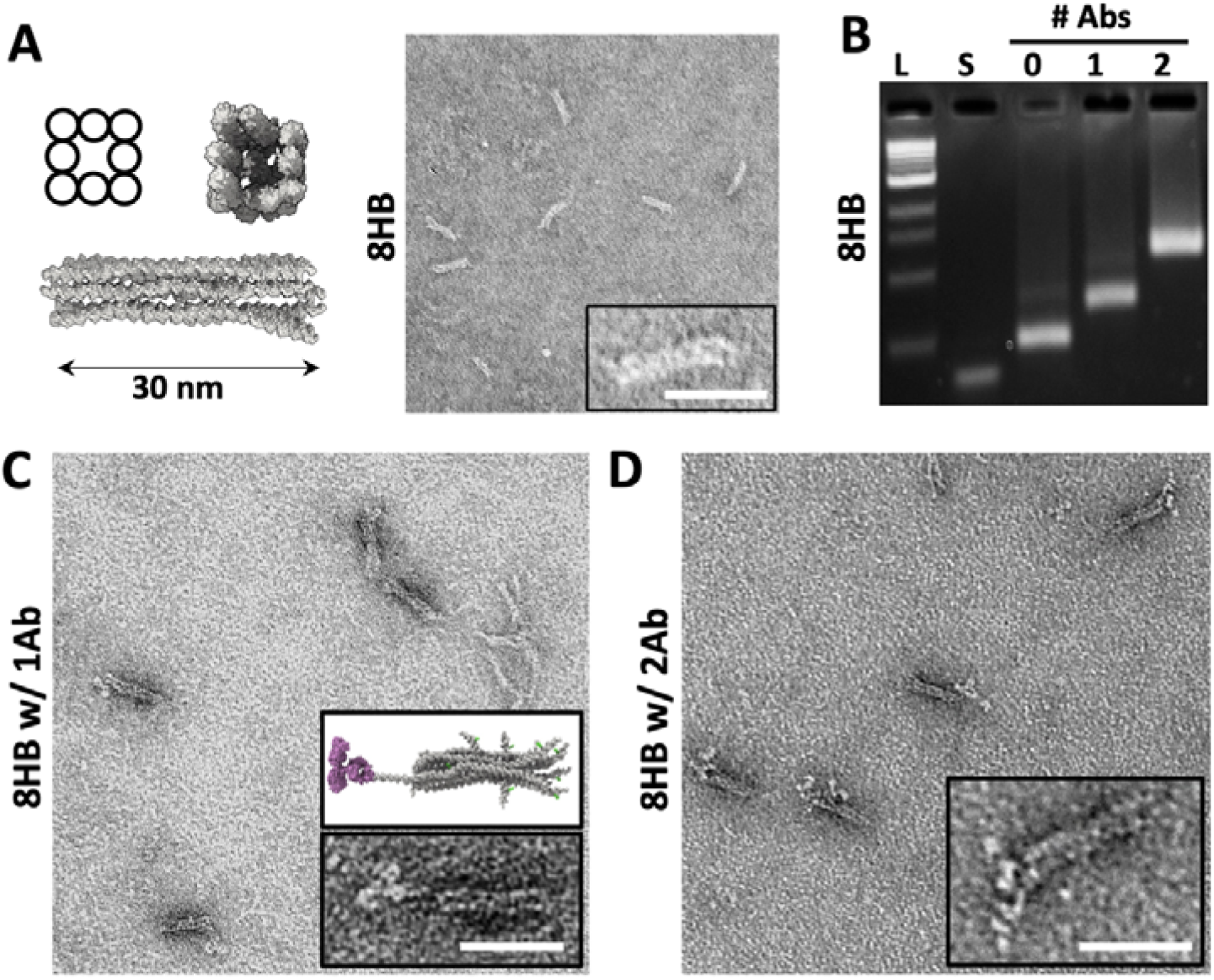
Fabrication and antibody labeling of DNA origami nanostructures. A) Design schematic, oxDNA simulation, and TEM image of the 8HB DNA origami structure. The simulation model depicts the base structure without the 8 overhangs for fluorophore attachment. B) Gel electrophoresis illustrates clear and efficient labeling of DO with one or two RNA pol-II Abs indicated by mobility shifts. TEM imaging confirmed efficient functionalization with 1 (C) or 2 (D) antibodies. Insets show a zoomed in depiction of a single functionalized DNA origami structure (compared to simulated 8HB structure with overhangs for fluorophore labels and antibody attached for size reference in (C)). Scale bars are 30 nm.

### Production of single stranded DNA (ssDNA) scaffolds

769-nt scaffold strands (sequence in Supplementary Table S2) were produced through PCR, using one 5’ phosphate modified to allow exonuclease digestion after PCR.[49], [50] The ssDNA scaffold was initially prepared using Guide-it Long ssDNA Production System v2 kit (Takara Bio, 632666) following manufacturer’s protocol. Subsequent larger-scale preparations were performed using PCR followed by lambda exonuclease digestion. Briefly, the target scaffold sequence was first amplified in double-stranded DNA (dsDNA) form via PCR from M13mp18 using PrimeSTAR Max Premix (Takara Bio, R045A) and 0.8 µM primers (Integrated DNA Technologies, primer sequences in Supplementary Table S3), where the reverse primer is modified with 5’ phosphate to facilitate selective exonuclease digestion. The PCR product was then mixed with 1/3 volume 10 M ammonium acetate (Sigma, A1542) and 2 volume ethanol (Sigma, E7023) to perform EtOH precipitation,[51] and the dsDNA pellet was resuspended in 5 mM Tris in ddH_2_O. To digest the anti-sense strand, dsDNA was mixed with lambda exonuclease (New England Biolabs, M0262) in the vendor supplied buffer, adjusted to 250 ng/μl DNA concentration, and then incubated at 37°C for 6 hr. 1 unit lambda exonuclease was added per 30µg DNA. After digestion, 10 mM EDTA was added to quench the reaction followed by heat inactivation at 75°C for 10 min. Digested ssDNA product was then mixed with 1 μl 20 mg/mL glycogen (Thermo Scientific, R0561) and 1/3 volume 10 M ammonium acetate to perform EtOH precipitation, and ssDNA pellet was resuspended in 1xTE buffer (10 mM Tris-HCl pH 8.0, 1 mM EDTA). Resuspended ssDNA was evaluated using gel electrophoresis on 1.5% agarose gel in 1xTAE buffer (40 mM Tris-base, 10 mM acetic acid, 1 mM EDTA) (Supplementary Figure S2).

### Folding, and purification of DNA origami nanostructures

DO nanostructures were folded according to established protocols.[52]–[54] Briefly, 20 nM scaffold ssDNA was mixed with a 10-fold excess of staple strands in Folding Buffer (FoB), (5 mM Tris, 1 mM EDTA, 5 mM NaCl, 20 mM MgCl_2_), and subjected to thermal annealing (BioRad C1000 Thermocycler). Details of the thermal annealing protocol can be found in the Supplementary Information (Supplementary Table S4). Agarose gel electrophoresis was used to evaluate the folding of the DO nanostructures. Agarose gels [2% agarose, 0.5× TAE with 10mM MgCl_2_, containing 0.5 μg/ml ethidium bromide] were run for 90 min at 90 V cooled in an ice-water bath or in a 4°C refrigerator. For nuclear delivery experiments structures were purified by centrifugation in the presence of polyethylene glycol (PEG)[55]. Briefly, the solution of folded DO was mixed with an equal volume of 15% PEG8000-based precipitation buffer, and spun at 16000g for 25 min to pellet DO. The pellet was resuspended in desired buffer to recover DO after discarding supernatant containing excess staple strands.

After purification through two rounds of centrifugal PEG precipitation, the DO was resuspended in 1x PBS with 2.5 mM MgCl_2_, and the concentration was measured using a Nanodrop (Thermo Scientific ND-ONEC-W). To label DO with fluorophores, the structures were designed to contain 8 ssDNA overhangs (i.e. staples that protrude from the bundle structure) to allow for binding a complementary oligonucleotide strands containing a Cy5 fluorophore label (sequences in Supplementary Table S1). Fluorophore labeled strands were designed to bind so the fluorophore is located near the surface of the structure. Fluorophore labeled strands were incubated with the structures at 20-fold molar excess with respect to the DO concentration. This excess corresponds to a 2.5 fold molar excess relative to the numbere of overhang strands on the origami structures. The mixture was then incubated at 37°C for 2 hr to allow for efficient binding of the fluorophore-labeled staples. The excess fluorophore-labeled overhangs were removed using a 0.5 ml 100 kDa MWCO Amicon filter unit by loading the sample into the filter unit (the total volume does not exceed the 0.5ml capacity of the filter) and centrifuging at 2000 g speed for 5 min. This filtration step was repeated 5 times with the addition of PBS buffer containing 2.5 mM MgCl_2_ buffer into the filter unit, which ensured the elimination of excess fluorophore labeled staple strands. The purified nanostructures were then stored at 4°C for subsequent antibody labeling.

### Antibody preparation

The mouse monoclonal antibody (mAb 7G5) specific for the C-terminal repeat domain (CTD) of the largest subunit of Pol II, RPB1-CTD (hereafter called anti-Pol-II antibody), and the mouse monoclonal antibody (17TF2-1H4) specific for the bacterial maltose-binding protein (anti-MBP, #MA3045 Fisher Scientific) were purified as described,[34] with minor modifications. MBP is not expressed in mammalian cells and hence provides a non-specific antibody control. Briefly, 1 ml antibody-containing ascites was incubated with 1.2 ml settled bead volume of pre-equilibrated Protein G Sepharose Fast Flow beads (GE Healthcare) for 2 hr at 4°C with gentle agitation. Beads were then transferred to a Poly-Prep Chromatography column (Bio-Rad) and washed for 20 column volumes with PBS. Antibodies were eluted in 1 ml fractions by 0.1 M glycine, pH 2.7, and were directly neutralized with 70-90 μl of 1 M Tris-HCl, pH 8.0. 6.5 µl aliquots from each fraction were analysed by SDS-PAGE and the fractions containing most of the antibodies were pooled and dialyzed in DiaEasy Dialyzer 6-8 kDa MWCO dialysis tubes (K1013-100, BioVision) against 2 liters of PBS overnight, and then for 2 hr with 2 liters fresh PBS. The antibody solution was then concentrated on Amicon Ultra-4 centrifugal filters with 10 or 50 kDa molecular weight cutoff (Millipore) to 1-4 mg/ml in PBS.

### Conjugation of antibodies with DNA

To initiate the process of DNA-antibody conjugation, 2 μl of DBCO-PEG5-TFP crosslinker (dissolved in dimethyl sulfoxide to a concentration of 900 µM) was combined with 1 mg/ml purified anti-Pol II antibody (or anti-MBP antibody) in 100 μl of PBS buffer (pH 7.4). The mixture was then incubated at 37°C with gentle shaking for 4 hr. Following this, the antibody-crosslinker product was removed through dialyzing for 3 time against 4 liters of PBS with a 6-8 kDa molecular weight cutoff dialysis membrane. The first two dialysis were done for 2 hours, while the third was done overnight.

Subsequently, the purified antibody-crosslinker product was combined with a 2-fold excess of azide-modified oligonucleotide (sequences in Supplementary Table S1) in PBS. The mixture was incubated at 37°C with gentle shaking for 2 hr, followed by incubation at room temperature overnight. To remove excess azide oligos, the sample was buffer exchanged 5 times into PBS with 2.5 mM MgCl_2_ with a 0.5 ml 100kD molecular weight cutoff Amicon filter unit. The resulting purified DNA-conjugated antibody was then stored at 4°C for DO functionalization.

### Functionalization of Antibody-labeled DNA origami

To conjugate DNA-labeled antibodies to DO, a solution containing 1 μM DNA-antibody conjugates was added to a solution of 100 nM fluorophore-labeled DO structures in a 100 μl buffer of 1x PBS containing 2.5 mM MgCl_2_. The mixture was thoroughly mixed and incubated at 37 °C with gentle shaking for 2 hr and then at room temperature overnight. The conjugation of the antibodies to the structures was confirmed with agarose gel electrophoresis (2% agarose, 0.5× TAE buffer, 10 mM MgCl_2_, and 0.5 μg/ml ethidium bromide) for 180 min at 90 V (Supplemental Figure S3). Samples for Transmission electron microscopy (TEM) imaging were purified using the Freeze ’N Squeeze (Bio-Rad) gel extraction column as per the manufacturer’s protocol. The target bands were excised from agarose gels, placed into the respective spin columns, and spun at 10,000 g for 5 min.

### Purification of Antibody-labeled DNA origami

All DO samples used in cellular experiments were purified via gel electrophoresis with electroelution, in order to obtain a pure product of DO labeled with zero, one, or two antibodies. The agarose gel (2% agarose, 0.5× TAE buffer, 10 mM MgCl_2_, and 0.5 μg/ml ethidium bromide) electrophoresis was done for a duration of 180 min at 90V while cooled in an ice bath or in a 4°C refrigerator. The desired bands were excised from the gel and placed in a dialysis membrane containing the same running buffer. The DO sample was then electroeluted from the gel fragment with a constant voltage of 90 V was applied for 1-2 hr until the product of interest had migrated out of the gel fragment and into the buffer as confirmed by the absence of ethidium bromide signal. The voltage was then reverse for 1-2 min to release any DO sample that was bound to the dialysis membrane. The DO sample was was recovered from the dialysis membrane with a syringe and filtered through a 0.2 μm filter to remove any remaining agarose. Finally, the DO sample was concentrated and buffer exchanged into 1x PBS with 2.5 mM MgCl_2_ using a Amicon filter with 100 kDa MWCO.

### TEM imaging of DNA origami structures

Samples for TEM imaging were prepared as previously described.[52], [56] Briefly, the DO sample was diluted to a concentration of 1-2 nM in 1x PBS containing 2.5 mM MgCl_2_. A glow-discharged copper grid was placed on a 10 μl drop of the DO sample on a parafilm sheet. The grid was incubated on the sample droplet for 4-6 min at room temperature to allow the DO structures to deposit onto the surface. Excess sample was removed by gently dabbing the edge of the grid with a piece of filter paper (Whatman). To stain the grid, two 10 μl drops of 2% uranyl formate (UFO) solution were deposited on a parafilm sheet. The first drop was applied onto the grid and immediately dried by gently dabbing the edge of the grid onto a piece of filter paper. The second drop was applied onto the grid and incubated for 5-10 seconds. Excess stain was then removed from the grid by again gently dabbing the edge of the grid with a piece of filter paper. The grid was allowed to dry for at least 20 min before imaging. TEM imaging was performed at the OSU Campus Microscopy and Imaging Facility on an FEI Tecnai G2 Spirit TEM using an acceleration voltage of 120 kV.

### Stability of DNA origami in cell culture media

Cy5-labeled DO samples were prepared at a concentration of 50 nM. Subsequently, 4 μl of each DO structure was mixed with 6 μl of the cell culture media (details of media provided in Cell Culture section). The mixture was incubated at 37°C for varying time periods (0, 1, 3, 6, 8, 12, 24 hr). To provide a baseline for comparison, a control sample of DO in 1x PBS with 2.5 μM MgCl_2_ buffer was prepared. For each time point, all samples, including the control, were evaluated by gel electrophoresis (2% agarose gel in 0.5x TAE buffer with 10 mM MgCl_2_ without ethidium bromide) run at 90V for 90 min cooled in an ice water bath or deli refrigerator. The resulting gel was imaged in a Cy5 channel, followed by post staining with 0.5 μg/ml ethidium bromide and imaging with UV excitation on a gel imager system (UVP GelStudio by Analytikjena). The integrity of DO structures of different sizes over time were assessed by comparing the gel electrophoretic mobility to their respective control sample. Additionally, high-resolution TEM images of the DO structure were to further evaluate the structural stability. The samples were purified for TEM imaging using the Freeze ’N Squeeze (Bio-Rad) gel extraction column and imaging samples were prepared as previously described.

### Stability of DNA origami in nuclear and cytoplasmic extracts

A volume of 4 μl of Cy5-labeled DO structures at 50 nM concentration was mixed with 6 μl of U2OS cytoplasmic and nuclear extracts both at 1 μg/μl (AscentGene) for a final DO concentration of 20 nM. The mixtures were incubated at 37°C for varying time periods (0, 1, 3, 6, 8, 12, 24 hr). After these incubations the structural integrity of the DO was evaluated using both gel electrophoresis (a 2% agarose gel in 0.5x TAE buffer with 10 mM MgCl_2_) run for 90 min at 90V cooled in an ice bath or at 4°C in a refrigerator. Structures were also evaluated by TEM to confirm structural stability following gel purifification using Freeze ’N Squeeze (Bio-Rad) gel extraction columns.

### Stability of DNA origami after electroporation

A volume of 4 μl of Cy5-labeled DO structures at 50 nM concentration were mixed with 6 μl R Buffer solution (Neon kits-MPK1096 Thermo Fisher). DO structures in R buffer were then immediately subjected to electroporation using the Neon Transfection system (MPK5000; Thermo Fisher) under the same conditions as cellular experiments using the 10-µl Neon tips with the following parameters: 1550 V, 3 pulses, and 10 ms per pulse. A control sample of each structure was also prepared in 1x PBS with 2.5 μM MgCl_2_ buffer for comparison and DO structures in R buffer solution but without electroporation were also examined as a control. The stability of the DO structures was characterized using gel electrophoresis and TEM imaging, as described above.

### Cell culture

A human osteosarcoma U2OS cell line (ATCC HTB-96) was obtained from the American Type Culture Collection (Manassas, VA). Cells were maintained in 5% CO2 atmosphere at 37°C in DMEM (4.5 g/L glucose) supplemented with 2mM GLUTAMAX-I, 10% FCS, 100 UI/ml penicillin and 100 μg/ml streptomycin.

### Electroporation of DNA origami into cells

Electroporation was performed using the Neon Transfection system with the 10-μl Neon electrode tips (MPK5000; MPK1096; Thermo Fisher) according to the manufacturer’s instructions. U2OS cells were washed once with Dulbecco’s phosphate buffered saline (DPBS, 14190136; Gibco) then trypsinized (0.25% Trypsin-EDTA; 25200-056; Gibco) for 3 min at 37 °C. Cells were then resuspended in R buffer to obtain 10 /10 µl cell suspension. 10 cells were mixed with 2 μl of 50 nM origami constructs and electoporated in the electrode tips using the following settings: 1550 V, 3 pulses, and 10 ms per pulse. The electroporated sample was then transferred directly into one well of an 8-well microscopy slide (Nunc Lab-Tek II for widefield imaging) containing 300 µl prewarmed medium without antibiotics and was incubated for 24 hr in 5% CO2 atmosphere at 37 °C.

### Sample fixation and Staining for Imaging of DNA origami in fixed cells

U2OS cells were fixed 24 hr after electroporation with 4% paraformaldehyde solution for 10 min at 37 °C, rinsed with DPBS, and subsequently incubated with (1:10,000) Hoechst 33342 trihydochloride, trihydrate (H3570; Invitrogen) stain solution in DPBS for an additional 10 min at room temperature and covered. Cells were rinsed again with DPBS and stored at 4 C until imaged.

### Fluorescence imaging of fixed cell samples

Cells were evaluated by HILO (highly inclined and laminated optical sheet) imaging with the Oxford Nanoimager S microscope (100x oil immersion objective, 1.45 NA, Hamamatsu Orca flash 4.0 CMOS camera, 200ms exposure time). The HILO illumination angle allows for widefield imaging within cell nuclei while reducing background. DPBS storage buffer was exchanged for an oxygen-scavenging imaging buffer (GLOX-14mg glucose oxidase, 20 mg/ml catalase, 10 mM Tris pH 8.0, 50 mM NaCl and 10% glucose in DPBS). Healthy, Hoechst-stained nuclei that expressed punctate Cy5 signal were identified and imaged sequentially (10 frames at 200ms exposure for each target) with excitation by 640nm laser (origami labeled with 8x Cy5) followed by 405nm (nuclei) in the same field of view and z-position. For two-color experiments (origami labeled with 4x Cy3 and 4x Cy5) cells were also imaged under the same exposure conditions under excitation with 560nm prior to the 405 nm.

### Confocal fluorescence imaging of fixed cell samples

Localization of 8HB DO in cells was also assessed by confocal microscopy. Fixed cells samples were imaged at 60x magnification using VT-iSIM high speed super resolution imaging system (VisiTech International) equipped with Olympus IX71 inverted, super-resolution VT-iSIM scan head, Hammamatsu ORCA Quest qCMOS camera, and 405, 442, 488, 514, 561, 640 nm excitation lasers. This system is best optimized for fast high-resolution confocal imaging of live or fixed samples. Image acquisition was set at the middle of the sample based on Hoeschst staining and Z-stacks of 5 μm of total thickness were acquired with 0.2 μm step size. Images were deconvoluted with microvolution plugin in ImageJ software.

### Processing of fixed cell images

Imaging data for all fixed cell experiments were processed using an in-house FIJI[57] macro. In brief, final images used for analysis were projections of the average fluorescence intensity over 10 separate frames of the same field of view (200 ms exposure). The image plane was selected by identifying the plane of largest nuclear area based on Hoechst signal.

### Analysis of nuclear delivery experiments

An in-house FIJI[57] macro was used to segment the nuclear region in images based on Hoechst signal and detect the fluorescent signal of origami within the segmented nuclear area. In brief, the Hoechst images were first segmented by Gaussian blur (sigma 8.0), then thresholded using Otsu’s method.[58] Thresholded nuclei were then made into regions of interest (ROIs) and their areas were measured. To quantify the origami signal, first a flat background fluorescence value was subtracted from all origami images based on the maximum background fluorescence found in control images in which no origami were present. Then, each nuclear ROI was used as a mask on its respective origami image to segment the origami signal within that nucleus. Origami structures with high enough fluorescence signal above the background were detected and counted using the Find Maxima function (prominence = 60) within these ROIs after application of Gaussian blur (sigma 4.0). For each nucleus, the origami count was divided by the nuclear area.

### Analysis of two-color delivery experiments

Colocalization of fluorescent signal in stability experiments was assessed with FIJI via JaCoP (Just another Colocalization Plugin).[59] In brief, fluorescent signal from origami labeled with Cy3, Cy5, or both fluorophores was thresholded using Otsu’s method[58], and the degree of colocalization between signals was calculated as the Pearson’s Correlation Coefficient.

### Live cell imaging of DNA origami in cells

For live cell imaging, 24 hr after electroporation, the medium of the electroporated U2OS cells on the 8-well microscopy slides was exchanged with 300 µl of fresh medium containing 500 ng/ml Hoechst33342 (H3570; Invitrogen) and no phenol red. After the medium change, cells were incubated for at least 10 min before imaging. The medium was then exchanged with 200 µl live-imaging compatible oxygen quenching buffer (DMEM without phenol red containing 22 mg/ml glucose, 67.3 mM HEPES [pH 8.0], 560 µg/ml glucose oxidase [G2133; Sigma], 40 µg/ml catalase [C1345; Sigma]), and cells were imaged immediately after adding the imaging buffer for no longer than 30 min on a HILO microscope.

HILO live cell imaging was performed on a home-built setup (system details previously described[60]) based on a NikonEclipse Ti microscope equipped with an acousto-optic tunable filter (AOTF; Opto-Electronic), a 100x 1.49 NA oil-immersion objective and a Hamamatsu EM-CCD camera (ImagEM X2 C9100-23B). Samples were imaged using a temperature-controlled on-stage chamber set to 37°C. The laser lines at 405 nm and 642 nm were used for excitation of Hoechst33342 and Cy5 fluorophores, respectively. Laser power during the experiments was set to 130 mW for 642 nm laser. Z-stabilization was ensured by the perfect focus system (PFS, Nikon Eclipse Ti) on the microscope. EM-CCD gain was set to 610, and samples were imaged with 10 ms exposure time over a total time of 10 s. After each time lapse, a single image of the nucleus based on Hoechst signal was also recorded in the 405 channel to identify the nuclear region.

### Live cell imaging data processing and analysis

Snapshots of live cell timelapses were generated and analysed in FIJI. First, maximun intensity projections of 5 consecutive frames of the Cy5 channel after the 20^th^ (200 ms) frames were generated for each time lapse. For the example images shown in Figure 5A, the Cy5 channel (shown in Red Hot LUT) was merged with the Hoechst image (shown in blue). For quantification (Figure 5B), the nuclei were segmented based on the Hoechst images, and the particles on the Cy5 images were detected inside the nuclear regions using the Detect Particle function of the ComDet v.0.5.5 plugin (https://github.com/UU-cellbiology/ComDet) with the following parameters: do not Include larger particles; do not Segment larger particles; Approximate particle size: 3 pixels, Intensity threshold (in SD): 5; ROIs shape: ovals. The detected particles were then visually curated and counted. For the statistical analyses a Kruskal-Wallis test with Dunn post-hoc analyses was sperformed in Python 3.11.5 (packages scipy and scikit_psthocs).

For tracking the motion of DNA origami structures, the live cell imaging sequences were first processed in FIJI[57] to correct for photobleaching. Plane correction from the BioVoxxel package[61] was used to flatten the signal. The Hoechst stain for each sample was used to determine the nuclear region, and the origami signal outside of the nucleus was removed. A custom CellProfiler pipeline was used to identify individual DO structures and obtain persistent particle tracks. Custom MATLAB scripts (codes are available at https://github.com/marcello-deluca/nuclear-origami-live-imaging-analysis) were used to calculate the diffusive behavior of the DO structures based on mean squared displacement (MSD):

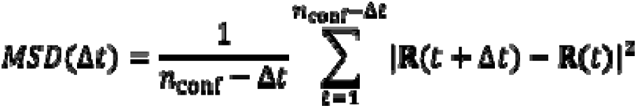

where □*t* is the quantity of elapsed time (expressed in frames), *n_conf_*is the overall number of configurations (frames) in a trace, *t* is a frame in the trace, and **R** is the 2D location of the origami at the specified frame of the trace determined in pixels and converted to nm based on a pixel size of 106.67 nm/pixel.

## RESULTS

### Design and fabrication of DNA origami nanostructures

To develop DO devices that are effective in cellular environment, we prioritized designs similar to structures previously shown to be stable in physiological conditions and resistant to degradation[36], [38]. We therefore focused on two rod structures with square lattice cross sections. The 8HB was designed in caDNAno[39], and the 26HB design has been previously reported[36]–[38]. Both the 8HB (∼6 nm x 6 nm cross-section and length of ∼30 nm, molecular weight of ∼0.5 MDa) and the 26HB origami structures (∼10 nm x 12 nm cross-section with a length of ∼90 nm, molecular weight of ∼5 MDa) were evaluated using coarse grained molecular dynamics simulations with the oxDNA model[45], [46] confirming a well-defined nanorod shape (Figure 2A and Supplementary Figure S4). The DO structures were faricated via molecular self-assembly and evaluated by gel electrophoresis and TEM (Figure 2A-B and Supplementary Figures S4-5). We leveraged the specific labeling capabilities of DO by adding ssDNA overhangs protruding from the structure to bind complementary strands with desired functionalities. The design included 8 side overhangs, specifically tailored for fluorophore labeling by binding to a complementary oligonucleotide labeled with a fluorescent Cy3 or Cy5 molecule. Overhangs for Cy3 and for Cy5 had distinct sequences to allow for attaching a defined number of each. Additionally, one or two ssDNA overhangs were included on one end of the DO to facilitate antibody attachment.

After the addition of fluorophore labeled strands and purification, the functionalized DO nanostructures were characterized by gel electrophoresis and TEM imaging (8HB shown in Figure 2, and 26HB shown in Supplementary Figure S4). Since the 26HB design was previoously reported and characterized[36], [38], here we focused our analysis on the 8HB. Gel electrophoresis revealed well-folded populations of 8HB structures with a single dominant population after purification by electroelution. Structures labeled with antibodies exhibited slower mobility with clear shifts between DO with 0, 1, and 2 antibodies with labeling yields of ∼70% or better (Supplementary Figure S3) with electroelution purification effectively removing unlabeled DO (Figure 2B). TEM imaging revealed well-folded nanorod-shaped structures about 30 nm in length, and one or two antibodies were visible for the single or double antibody-labeled designs (Figure 2C-D, Supplementary Figure S5). The insets in Figure 2C-D show zoomed in TEM images of the labeled 8HB, and for the single antibody label a simulated version is also shown for comparison. We only studied the effects of antibody number on the 8HB, so the 26HB was only labeled with a single antibody (Supplementary Figure S4). These findings highlight the precision of the design and emphasize the controlled assembly and specific labeling capabilities of the structure.

### Stability of DNA origami nanostructures in biological solutions

To confirm the suitability of the fabricated DO structures for intracellular applications, we first tested the stability of the structures in multiple relevant biological solutions including cell culture media and cellular cytoplasmic and nuclear extracts prepared from human cells (U2OS). We monitored the structural integrity of DO over a 24 hr period using agarose gel electrophoresis, imaging gels in the Cy5 channel to confirm stability of overhang attachment. Hereafter we refer to the DO with fluorophores added to the 8 attachment sites as 8HB-8Cy5 or 26HB-8Cy5. Gel analysis revealed structures consistently showed the same gel mobility and intensity, indicating structural integrity and stability in cell media (8HB-8Cy5 results in Figure 3A and Supplementary Figure S6, and 26HB-8Cy5 results in Supplementary Figure S7), which is consistent with prior studies on the 26HB design[38]. This analysis was extended to extracts to consider the stability inside cells, revealing that the 8HB-8Cy5 and 26HB-8Cy5 remained highly stable throughout the 24 hr monitoring period in U2OS cytoplasmic and nuclear extracts (8HB-8Cy5 results in Figure 3B-C and Supplementary Figure S8, and 26HB-8Cy5 results in Supplementary Figure S9). For TEM imaging, we focused on the 24 hr timepoint in the nuclear extract as the most relevant condition. TEM analysis further demonstrated intact structures in nuclear extract at the 24 hr timepoint (Figure 3C, and Supplemental Figures S8-S9). While it is challenging to accurately recapitulate an intracellular environment, these results suggest these DOs can remain stable over extended times in the presence of cytoplasmic and nuclear components. Hence, these results indicate DO nanostructures could be well-suited for intracellular applications that require the structure to remain intact, although the integrity of the structure once introduced directly inside live cells is still important to verify, which we address subsequently.

**Figure 3:**
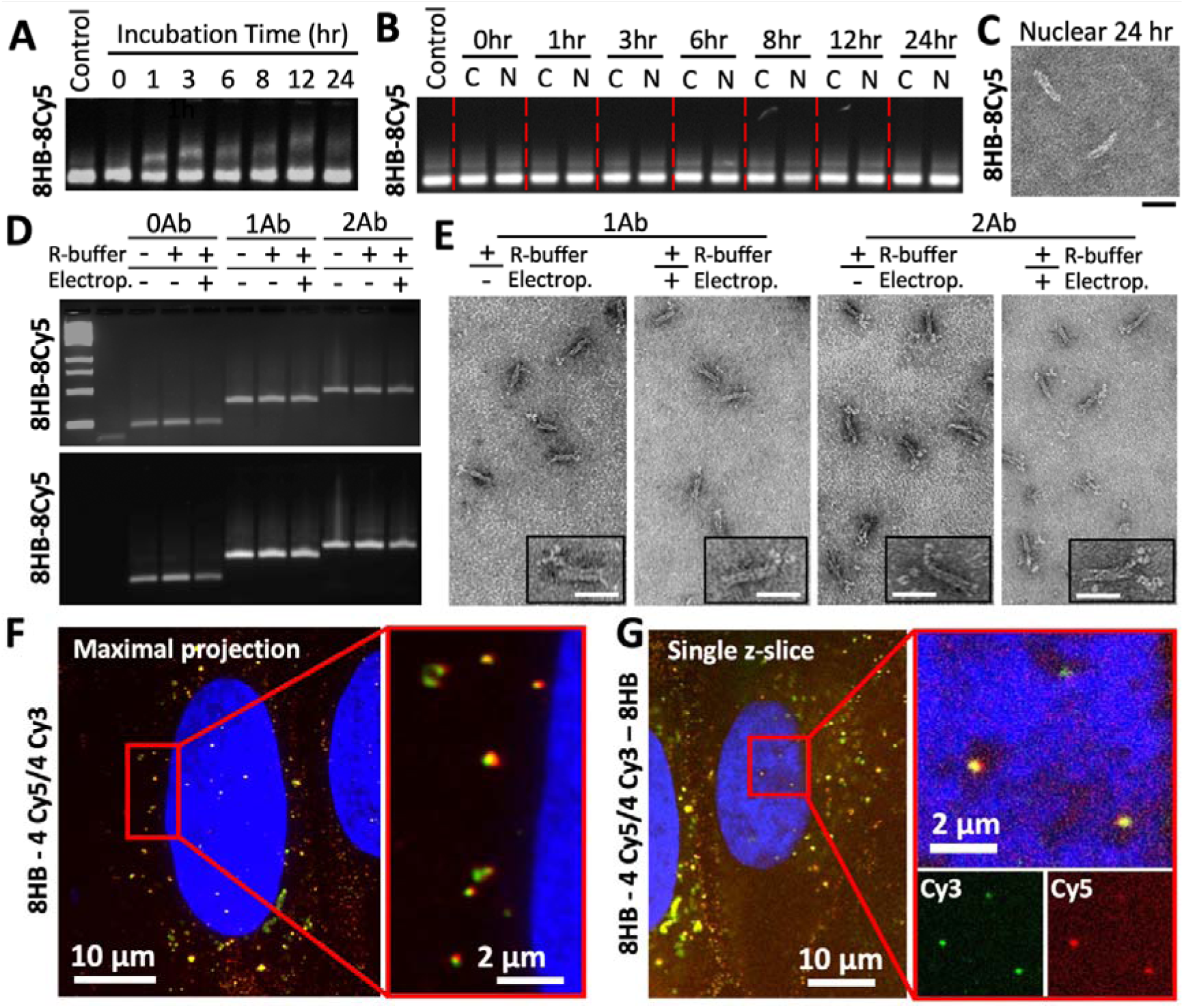
Stability of DNA origami nanstructures. A-B) Agarose gel electrophoresis (images show Cy5 fluorophore emission) revealed consistent mobility when 8HB structures were incubated at 37 C in A) cell culture media or B) cytoplasmic (C) or nuclear (N) extract from U2OS cells, verifying structural stability and fluorophore attachment integrity up to 24 hr. C) TEM imaging also confirmed structures remain intact, shown for the 24hr nuclear extract condition (Scale bar is 30 nm). D) Agarose gel electrophoresis also revealed no changes in mobility in electroporation buffer (R-buffer) and after being subjected to electroporation for structures alone or for structures with 1 or 2 antibodies (Ab) attached. The Ethidium Bromide stain is shown on top and Cy5 emission channel on bottom. E) TEM imaging also confirmed structures remain intact and antibodies remain attached in R-buffer and after electroporation. Insets show zoomed in views of a single 8HB structure with 1 or 2 antibodies attached (Scale bars are 30 nm). F) and G) 8HB double-labeled with Cy3 and Cy5 electroporated into U2OS cells exhibit co-localization of both fluorophores in the cytoplasm (F) and the nucleus (G). Scale bars are indicated in the panels and insets.

### Electroporation of DO structures into U2OS cells

We chose electroporation as a mechanism to get DO structures into cells, which has previously been demonstrated as an effective mechanism to deliver gene encoding DOs into live cells[14], [15], [32]. To test the viability of electroporation for the delivery of intact DO into nuclei, we first performed electroporation experiments with Cy5-labeled 8HB and 26HB DO with no antibodies attached. These initial electroporation tests revealed the 26HB exhibited significant aggregation when introduced into U2OS cells (Supplemental Figure S10). We attribute this to the size of the 26HB (approximately 4.8 MDa or 7,200 bp in total size); prior studies have shown similar aggregation behavior of other electroporated nanomaterials such as quantum dots[62], silver nanoparticles[63], and DNA plasmids[64]. On the other hand, the 8HB exhibited minimal aggregation and distributed more homogeneously throughout the cell cytoplasm. Hence we focused the remaining experiments on the 8HB (Supplemental Figure S10).

### Stability of DNA origami nanostructures after electroporation

We next performed experiments to assess the stability of antibody labeled 8HB before and after electroporation using gel electrophoresis and TEM. These experiments were carried out with DO labeled with anti-Pol-II antibodies. As a control, we also tested the stability of DO in the manufacturer provided R buffer, used for cell resuspension before electroporation. Gel electrophoresis revealed that the structures (unlabeled and labeled with one or two anti-Pol II antibodies) exhibited similar mobility in R buffer before and after electroporation compared to a control structure kept in storage buffer and not subjected to electroporation (Figure 3D). Gels were imaged both in the Cy5 channel and with ethidium bromide staining of the DNA, suggesting that the DO structure and the fluorophore and antibody labeling all remained intact after electroporation. TEM imaging of gel-purified samples confirmed that DNA origami structures maintained their shape and antibody labeling in R buffer and after electroporation (Figures 3E).

### Stability of DNA origami nanostructures in U2OS cells

To directly assess the stability of fluorophore-labeled nanostructures after transfection into cells, we performed electroporation experiments with DO labeled with two distinct color fluorophores Cy3 and Cy5 (i.e. double-labeled structures). We reasoned that if DO structures remain intact, the Cy3 and Cy5 emission would remained co-localized. Importantly, prior work has shown that even degradation products of DNA structures can exhibit fluorescence co-localization in cells.[65] In addition, aggregation of DO structures can also lead to the apperance of co-localization. To account for these possibilities, we performed experiments in which 8HB structures were either double labeled with Cy3 and Cy5, or singly labeled with either Cy3 or Cy5, but co-electroportated into cells. If the 8HB structures and fluorophore labeling are stable, we predicted that the colocalization between Cy3 and Cy5 signal in double-labeled 8HB structures would be higher than in the co-electroporation with the single-color structures. On the other hand, if the 8HB structures were unstable (i.e. subject to intracellular degradation) or aggregated, we expected colocalization values similar to the co-transfection condition.

U2OS cells were electroporated with 8HB structures in both experimental configurations: 8HB structures double-labeled with Cy3 and Cy5, and a co-transfection of 8HB labeled with Cy3 and 8HB labeled with Cy5. U2OS cells were fixed and nuclei stained 24 hr after electroporation. Cells were imaged with HILO illumination and the nuclear stain was used to set the focus to the mid-plane of the nucleus. In addition, we carried out iSIM (instant structured illumination microscopy) imaging in which z-stacks were recorded throughout the volume of the nucleus. We observed clear visual co-localization of Cy3 and Cy5 signals in the double-labeled condition in both HILO and iSIM images, while co-transfection gave rise to visually lower levels of co-localization visualized via HILO imaging (Figure 3F-G and Supplemental Figure S11). To quantify the spatial correlation between the two signals, we calculated the average Pearson correlation coefficient, *r*, for the detected Cy3 and Cy5 fluorescence peaks in HILO images. The 8HB structures exhibited correlation coefficients of r = 0.82 ± 0.10 for the double-labeled 8HB (8HB-4Cy5/4Cy3) and r = 0.34 ± 0.14 for the co-transfected single labeled 8HB (8HB-8Cy5 plus 8HB-8Cy3). These results suggest the 8HB DO remain primarily intact inside cells up to 24 h after electroporation. Importantly, in the iSIM images, we observed 8HB DO inside the nucleus that exhibited co-localization, indicating DO are structurally stable at 24 hr even after entering the nucleus (Figure 3G, and Supplementary Figure S12). These results confirm the stability of 8HB in the cytoplasm and reveal that structures delivered into the nucleus also remain intact.

Combined, our results indicate that the 8HB DO nanostructure is structurally stable after electroporation, for at least 24 hr when exposed *in vitro* to cytoplasmic and nuclear extracts, and remains stable for 24 hr after electroporation into U2OS cells both in the cytoplasm and in the nucleus.

### Fluorescence imaging of DNA origami in fixed U2OS cells

To assess the efficiency of the piggybacking approach via Pol II antibodies as a delivery method of fluorescently labeled DO to cell nuclei, U2OS cells were electroporated with 8HB-8Cy5 conjugated to either zero antibodies (hereafter called 8HB-8Cy5), one anti-Pol II antibody (hereafter called anti-Pol II 8HB-8Cy5 1Ab), or two anti-Pol II antibodies (hereafter called anti-Pol II-8HB-8Cy5 2Ab). We also tested 8HB-8Cy5 structures conjugated with one anti-MBP antibody that has no endogenous targets in human cells (hereafter called anti-MBP 8HB-8Cy5 1Ab) and 8HB with no Cy5 and no antibodies (hereafter called 8HB-No Cy5) as controls. U2OS cells were fixed and nuclei stained 24 hr after electroporation. Cells were imaged using HILO illumination. Hoechst nuclear stain was used to locate and focus on the mid-plane of the nucleus to visualize DO within the nuclear interior. The individual DO structures appeared as bright, diffraction limited spots throughout the cytoplasm and within the nucleus (Fig 4A), which were absent in negative controls in which cells were electroporated with buffer alone or unlabeled DO (Supplemental Figure S13). In addition to diffraction limited punctate structures, we also observed large and bright spots mainly located within the cytoplasm (Fig. 4A), which likely correspond to aggregated DO structures. To determine the number of structures within the nucleus, we employed a custom Fiji macro, which used the nuclear stain as a mask. Individual spots that fell within the mask and had intensity above a threshold value (see Materials and Methods) were counted as a DO particle, likely corresponding to an individual structure. Brighter spots could correspond to multiple structures in close proximity, but these were still counted as a single particle in our analysis. The number of particles was normalized by the nuclear area to determine the density of DO within each nucleus at the midplane (Figure 4B). We found that the conjugation of 1 or 2 RNA Pol II antibodies to DO increased the overall number of 8HB-8Cy5 structures delivered to cell nuclei when compared to unconjugated DO or DO labeled with one anti-MBP antibody (Fig. 4B). The quantified densities correspond on average to 22, 105, and 96 DO particles at the nuclear midplane for the 8HB with 0, 1, or 2 anti-Pol II antibodies, respectively. These results were further confirmed via confocal imaging in which antibody conjugated 8HB-8Cy5 were detected inside the nuclei in single z-slices (Supplementary Figure S14).

**Figure 4:**
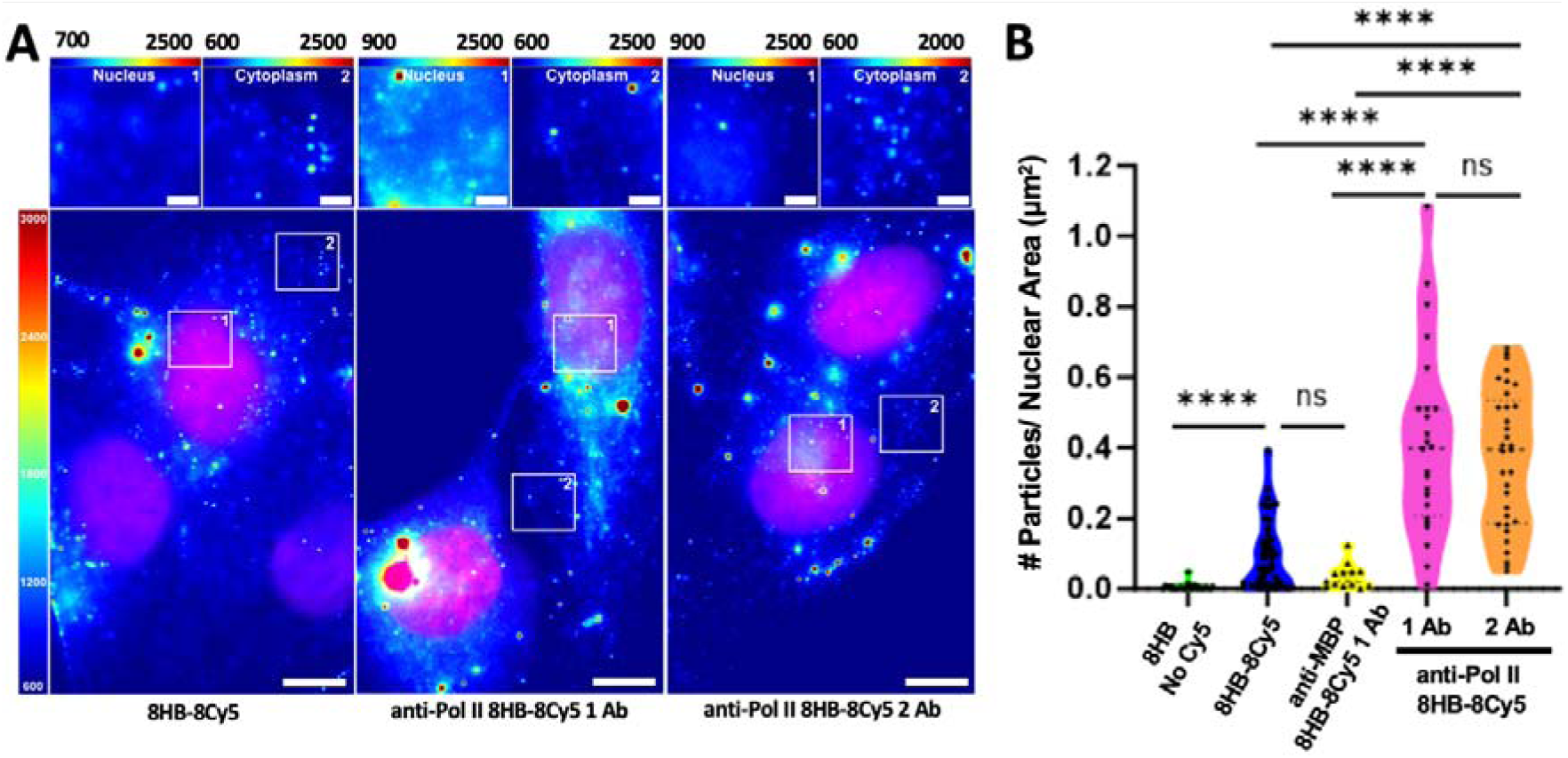
Pol-II antibodies facilitate piggybacking DNA origami to the nucleus. A) HILO imaging at the mid-plane of U2OS cells illustrates DNA origami structures inside cells for 8HB-8Cy5 with 0 antibodies (left, 8HB-8Cy5), 1 Pol-II antibody (middle, anti-Pol II 8HB-8Cy5 1 Ab), or 2 Pol-II antibodies (right, anti-Pol II 8HB-8Cy5 2 Ab). A clear increase in Cy5 fluorescence emission in cell nuclei is evident when 8HB-8Cy5 are labeld with 1 or 2 Pol-II antibodies. Upper images show a zoomed in views of the nucleus and the cytoplasm for each condition. Scale bars are 10 μm. B) The number of observed particles in the nuclei was quantified for each condition, showing some 8HB-8Cy5 enter the nucleus even without antibodies, and there is a significant increase in nuclear localization with 1 or 2 antibodies on 8HB-8Cy5 structures.

**Figure 5:**
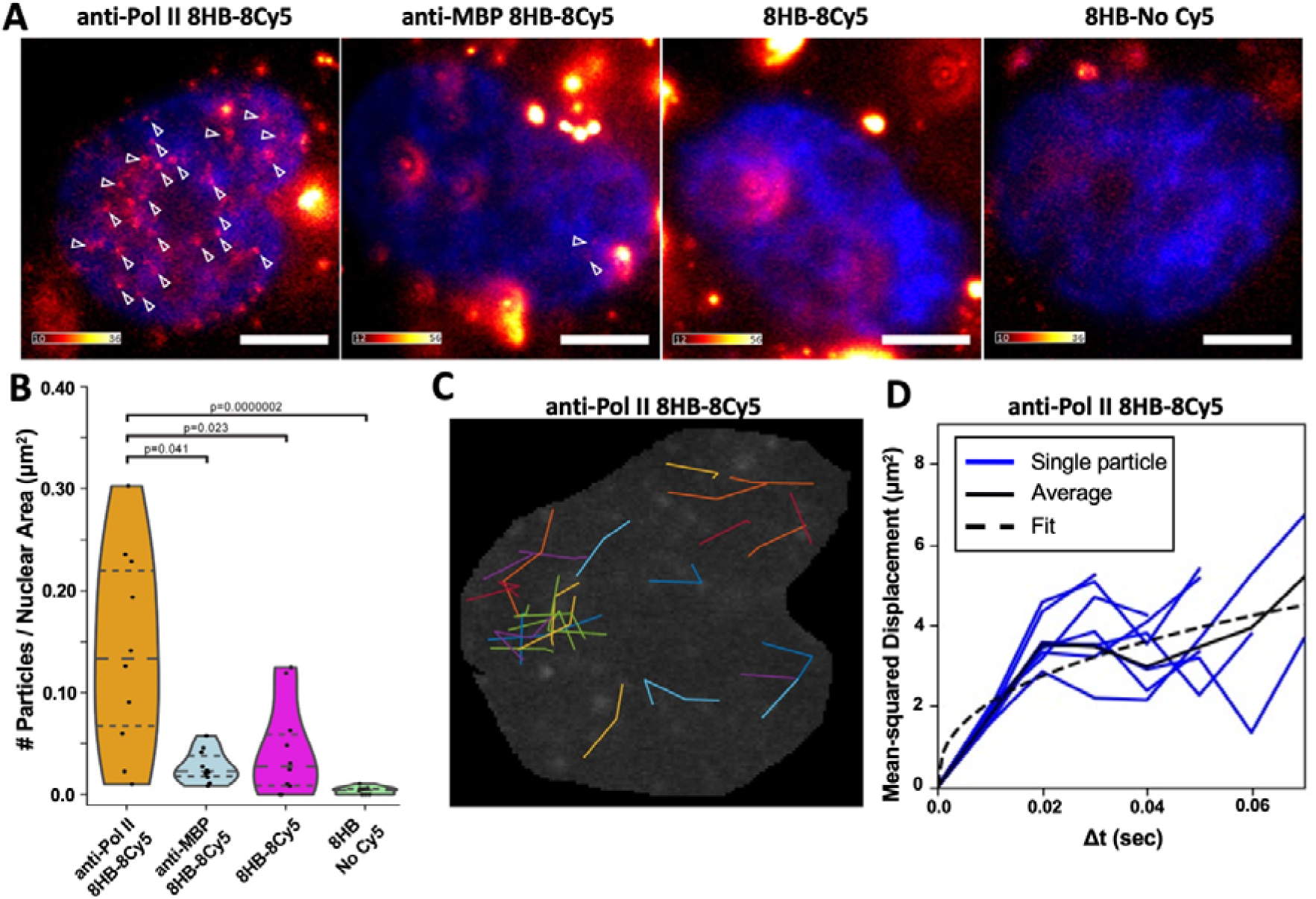
Nuclear delivery of DNA origami in live cells. A) Representative images of live U2OS cells 24 h after electroporation with non-functionalized 8HB DO structures (right, 8HB No Cy5) or 8HB-8Cy5 DO structures functionalized with either one anti-Pol II antibody (anti-Pol II 8HB-8Cy5; left), or one anti-MBP antibody (anti-MBP 8HB-8Cy5; 2nd from left), or 8HB-8Cy5 with no antibodies (8HB-8Cy5; 2nd from right). White arrowheads point to nuclear particles representing single DNA origami structures. Color bars indicate the fluorescence intensity range of the Cy5 signal (scale is x1000). Nuclear Hoechst staining is shown in blue. Scale bars are 5 µm. B) Combined violin-, box- and jitter-plots showing the quantification of nuclear particles. A significantly higher number of particles per nuclear area were detected in cells that were electroporated with anti-Pol II 8HB-8Cy5 compared to the other three conditions (H=22.6447, p=4.789x10^-5^, Kruskal-Wallis test). p values for pairwise comparisons using Dunn post-hoc analyses are shown. n=10. C) Traces of tracked particles for anti-Pol II-8HB-8Cy5 within the nucleus depicted in 5A. D) Average MSD data measured from each of the 7 nuclei analyzed and power-law fit over the entire data showing anomalous diffusion with an exponent of 0.4 (95% CI 0.1-0.7).

### Visualizing delivery of DNA origami nanostructures into live cell nuclei

To demonstrate the RNA Pol II-facilitated nuclear delivery of DO nanostructures in living human cells we electroporated anti-Pol II antibody functionalized 8HB-8Cy5 structures into U2OS cells. Since our prior results revealed no significant benefit to the incorporation of two anti-Pol II antibodies, we only performed live cell experiments with the single antibody-conjugated DO (labelled as anti-Pol II 8HB-8Cy5 in Figure 5). As controls, we also tested 8HB-8Cy5 linked with one anti-MBP (labeled as anti-MBP 8HB-8Cy5 in Figure 5), as well as 8HB-8Cy5 with no antibodies (labeled as 8HB-8Cy5 in Figure 5), or 8HB structures without any antibodies or Cy5 (labeled as 8HB-No Cy5 in Figure 5). 24 hr after electroporation nuclei were stained with Hoechst, and cells were imaged using live HILO microscopy with high temporal resolution (100 fps) for 10 seconds (Supplemetary Videos S1). We first compared the number of 8HB-8Cy5 structures measured in the nucleus of live cells as observed from the nuclear Cy5 signals. To identify these nuclear signals, we used maximum intensity projected images of 5 consecutive frames from the 210-250 ms timepoints from each sample. This revealed numerous diffraction-limited spots that appeared highly abundant in case of the anti-Pol II 8HB-8Cy5 sample. Quantification of the number of particles revealed significant enrichment in cells electroporated with anti-Pol II 8HB-8Cy5 compared to all three control conditions (Figure 5B). These results are consistent with our fixed cell imaging results and show that 8HB DO structures functionalized with RNA Pol II-specific antibodies are targeted to the nuclei of living human U2OS cells.

Timelapse imaging revealed clear motion of the anti-Pol II 8HB-8Cy5 structures inside the nucleus as shown by the time lapse recordings (Supplementary Video S1). Comparing the anti-Pol II 8HB-8Cy5 to either the anti-MBP 8HB-8Cy5, the 8HB-8Cy5, or the 8HB no Cy5 revealed that mobile 8HB DO structures were not, or hardly, visible in the control cases (Figure 5A and Supplementary Video S1-S2). For the anti-Pol II 8HB-8Cy5 case, the motion of individual structures could be tracked over multiple frames (10 ms per frame) while the structures remained in the image plane. We tracked the motion of 161 individual anti-Pol II 8HB-8Cy5 structures (several example trajectories are shown in Figure 5C). The length of these trajectories was limited to only a few tens of milliseconds due to the movement of particles out of the image plane. Nevertheless, calculating the mean squared displacement (MSD) over these short imaging periods as a function of time revealed that structures exhibit anomolous diffusion in nuclei following a power law behavior, *MSD* ∝ t , with a coefficient of approximately 0.4 (95% CI 0.1-0.7),. This coefficient of < 1 indicates sub-diffusive motion likely due to the crowded and viscoelastic nuclear environment, which is consistent with the sub-diffusive motion of other nuclear factors like the transcription factor P-TEFb, which was previously reported to exhibit a power law coefficient of = 0.6 inside the nucleus[66].

## DISCUSSION

DO nanostructures have been demonstrated for applications like biophysical measurements[8], [67], manipulating molecular interactions[9], [10], and delivery of therapeutic agents[2], [7], [68], which could all be useful intracellular functions; and other applications like high resolution imaging[22], nucleic acid and protein detection[2], [69], probing of chromatin sub-structures[28], [29], and gene delivery[14]–[16] could particularly benefit from mechanisms to specifically deliver DO to live cell nuclei. As a critical step for intracellular delivery and applications, we evaluated the stability of DOs in relevant conditions including cell culture media, cell ctyoplasmic and nuclear extracts, upon electroporation, and inside cells. Our results show that the DO designs used here are stable in cell media and in nuclear and cytoplasmic extracts for 24 hr, which is consistent with prior work showing DO can exhibit extended stability in cell culture or in cell lysates[36], [70], [71]. It is worth noting the stability is design dependent and DO can degrade more rapidly at higher serum levels[72], [73], which is an important consideration especially for translational applications. However, multiple strategies exist such as UV cross-linking or polymer coating and brushes that can extend the stability of DOs [74]–[76]. Prior work has shown that the process of electroporation can impact structural integrity of DO [77], while others studies have demonstrated some DO designs can remain stable through electroporation [14], [15], suggesting the electroporation stability is dependent on the design and electroporation parameters. Our results show that the 8HB DO structure and antibody attachment is stable after electroporation. We also demonstrate that the 8HB DO can remain stable for 24 hr after electroporation into cells in the cytoplasm or after entering the nucleus. While prior work have not evaluated DO inside nuclei, our results are in agreement with prior studies showing some DNA nanostructure designs can exhibit extended stability inside cells [78]–[80].

Several prior efforts have studied interactions between DO and cells (e.g. see recent reviews[12], [81]), and a few recent studies have demonstrated effective delivery of gene sequences folded into DO structures where genes can be expressed[14]–[16], [32]. Two of these studies leveraged either Cas9[14] or an SV40 derived DNA sequence[32] to promote delivery to the nucleus. However, these studies were focused on delivering information through the DNA sequence to the nucleus, rather than intact DO structures. Unlocking potential device functions of DO inside cell nuclei requires methods that allow for the delivery and tracking of intact DO into live cells and targeted delivery to nuclei. Here we targeted DOs to the nucleus by functionalizing them to bind neosynthetized nuclear factors in the cytoplasm, in this case the largest subunit of the RNA polymerase Pol II. As the nuclear factor is imported to the nucleus, the DO can be carried, or “piggybacked,” along with them.

We found this piggybacking approach is size dependent, with no clear nuclear delivery observed using a larger size DNA origami (∼4.8 MDa, ∼90 nm long nanorod), while the piggybacking approach worked effectively to deliver smaller structures (∼0.5 MDa, ∼30 nm long nanorod) to the nucleus. We confirmed that these DO remain intact inside cells for 24 hr using two-color fluorescence co-localization (DO dual-labeled with Cy3 and Cy5), including comparison to co-delivery of a single-labeled structures (Cy3-labeled DO plus Cy5-labeled DO) to verify that co-localization is the result of intact structures[65]. iSIM imaging further revealed DO can remain intact in live cell nuclei 24 hr after electroporation, hence opening a door to leverage the diverse functions of DO inside the nucleus. These ∼30 nm nanorod DO already provides a useful basis for functions like imaging, or detection with the simple inclusion of fluorophores or aptamers[82]. Our results further showed these DO are mobile inside the nucleus. They exhibit sub-diffusive motion similar to what has previously been measured for other nuclear factors[66], which is likely due to the highly constrained environment inside the nucleus. Nevertheless, our results suggest the piggybacked DO can explore the nuclear volume.

Some functions of DO would likely be enhanced through the use of larger structures. Here a key factor limiting our use of the larger 26HB DO was aggregation in the cytoplasm. The large design space of DO in terms of size, shape, surface coating, and functionalization can likely enable engineering of intracellular behaviors like aggregation, passive or active transport, and entry to the nucleus or other cell compartments. Our results and other recent efforts[12], [14], [32], [65], [78], [80] provide a framework to guide these studies. In the future, a better understanding of these intracellullar behaviors of DO will be important to for enabling additional applications, for example those that leverage multi-component devices like biophysical measurements[28].

The piggybacking approach we presented here relies on binding neosynthetized nuclear factors in the cytoplasm that will be imported to the nucleus. Here we targeted the RNA polymerase II building on prior studies that established the piggybacking approach for delivering antibodies to the nucleus.[33]–[35] These studies used the same approach to target multiple transcription factors, including TATA binding protein (TBP), TBP-associated factor 10 (TAF10), suggesting these, and likely a variety of other nuclear factors, could be used to piggyback DO structures to the nucleus. These proteins have specific mechanisms that drive localization to the nucleus, such as interactions with other proteins (e.g. RNA Pol II associated protein, RPAP2[83]) that mediate trafficking or direct interactions with importins via nuclear localization signal (NLS) sequences or other domains[84]. Indeed, prior studies showed the expression of gene sequences delivered via DO is increased with inclusion of either amino acid NLS or DNA nuclear targeting sequences (DTS)[14], [32]. Combined with our results, these studies suggest a variety of proteins or motifs or direct inclusion of NLS or DTS sequences onto DO could be alternative routes to specifically deliver intact DO devices to the nucleus.

### Authorship contribution statement

GM led all preparation of functionalized and purified DNA origami and performed gel electrophoresis and electron microscopy imaging. PC performed fixed cell HILO imaging experiments and analysis. AO performed antibody purification, electroporations, live cell HILO microscopy experiments, and data analysis. ES performed electroporations and confocal and iSIM imaging. WP designed the DNA origami structures, performed simulationof some DNA origami designs, and supported electron microscopy analysis. SS supported functionalizatioin and purification of DNA origami structures. YW supported the fabrication, gel electrophoresis, and electron microscopy characterization of DNA origami. PD supported live cell HILO microscopy and provided instrumentation live cell for imaging. MD performed simulation of some DNA origami designs and performed the particle tracking analysis for live cell experiments. GA supervised the simulation and particle tracking analysis. LT supervised the antibody preparation, live cell imaging experiments, and many aspects of the cellular work; and he provided funding and facilities. ML supervised all fixed cell imaging experiments and provided general guidance for imaging work; and she provided funding and facilities. MP supervised functionalization and purification of DNA origami and co-supervised characterization of DNA origami; and he provided funding and facilities. CC supervised the DNA origami design and fabrication and co-supervised DNA origami characterization; and he provided funding and facilities and led the overall collaborative project. GM, PC, AO and CC led the drafting of the manuscript with extensive input and feedback from all authors.

## Acknowledgements

We thank the present and past members of the Castro, Poirier, Lakadamyali, Tora, and Arya labs for insightful discussions. We thank the University of Pennsylvania, Cell and Developmental Biology Core Microscopy Facility and Dr. Andrea Stout for help with iSIM imaging. We also acknowledge support from the Campus Microscopy and Imaging Facility (CMIF) at The Ohio State University for TEM imaging. This work was financially supported by NSF-EFRI (Award Number:1933344) grant to MPG, LT, ML, GA, and CC; ANR-22-CE11-0013-01_ACT; Fondation pour la Recherche Médicale (EQU-2021-03012631) grants to LT; NIH MIRA (R35GM139564) grant to MGP and LT. This work, as part of the ITI 2021-2028 program of the University of Strasbourg, was also supported by IdEx Unistra (ANR-10-IDEX-0002), and by SFRI-STRAT’US project (ANR 20-SFRI-0012) and EUR IMCBio (ANR-17-EURE-0023) under the framework of the French Investments for the Future Program (for LT).

