## Supplementary Figure for "Piggybacking functionalized DNA nanostructures into live cell nuclei"

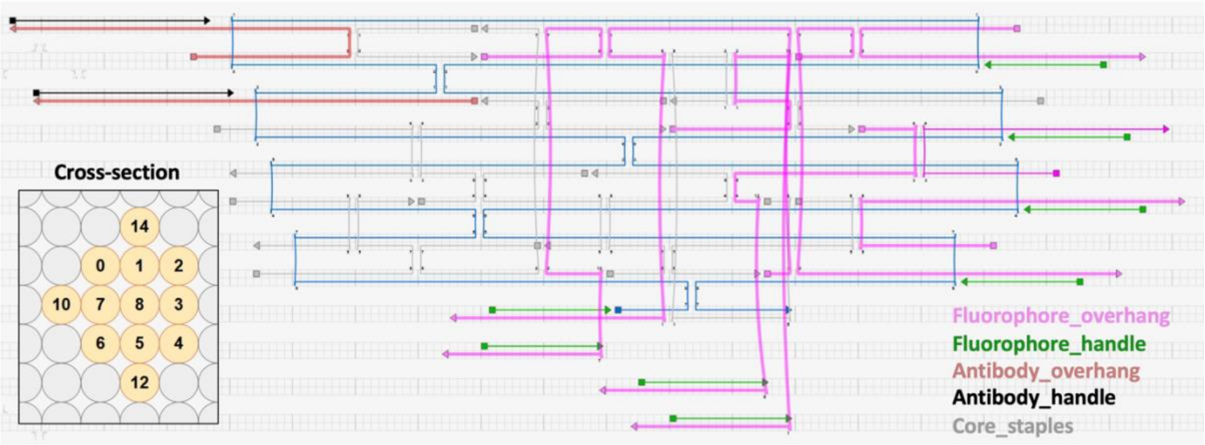


**Figure S1:** Cadnano design of 8HB. The bottom left inset shows a cross-section view. Helices 0-7 form the main structure of the 8HB. Helices 10, 12, and 14 are used for placing overhangs for fluorophore and antibody attachment as indicated by the color scheme.

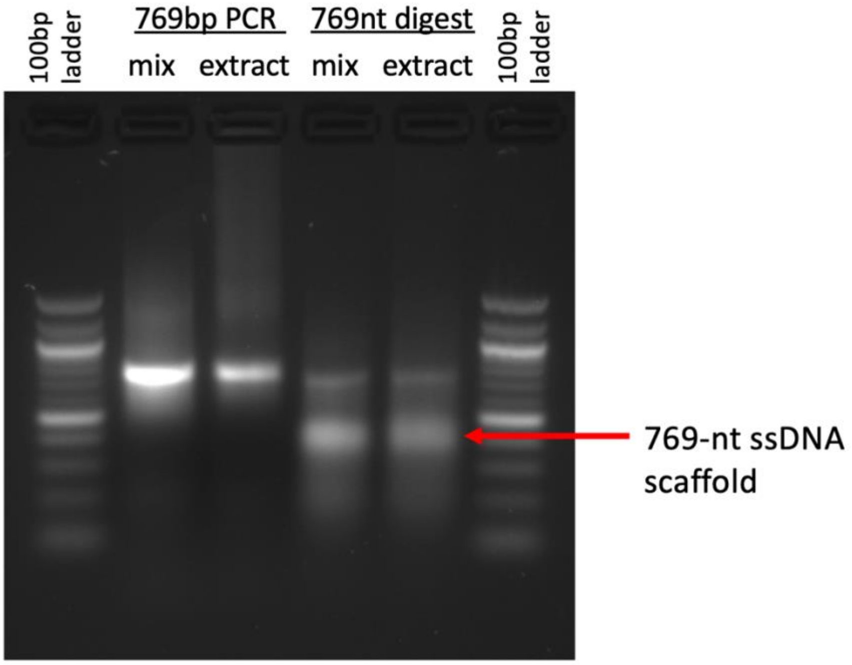


**Figure S2:** Agarose gel electrophoresis analysis of single-stranded DNA (ssDNA) scaffold preparation for 8HB folding. The lanes show a 100 base-pair (bp) ladder (lane 1); the 769 bp PCR product before (lane 2) and after purification by ethanol precipitation (lane 3); the 769 nucleotide (nt) ssDNA scaffold after lambda exonuclease digestion the phosphate labeled strand in the PCR product before (lane 4) and after purification by ethanol precipitation (lane 5); and again 100 bp ladder (lane 6). The red arrow highlights the 769 nucleotide ssDNA scaffold product.

**Table S4:** Steps of thermal annealing protocol used to fold DNA origami

| Temp [°C] | time [min] |
|---|---|
| 65 | 15 |
| 64-61 | 3 |
| 60 | 5 |
| 59-57 | 10 |
| 56 | 25 |
| 55 | 30 |
| 54 | 45 |
| 53-49 | 60 |
| 48-45 | 42 |
| 44 | 36 |
| 43-42 | 32 |
| 41-39 | 20 |
| 38 | 15 |
| 37 | 10 |
| 36-35 | 5 |
| 34-30 | 2 |
| 20 | Hold |

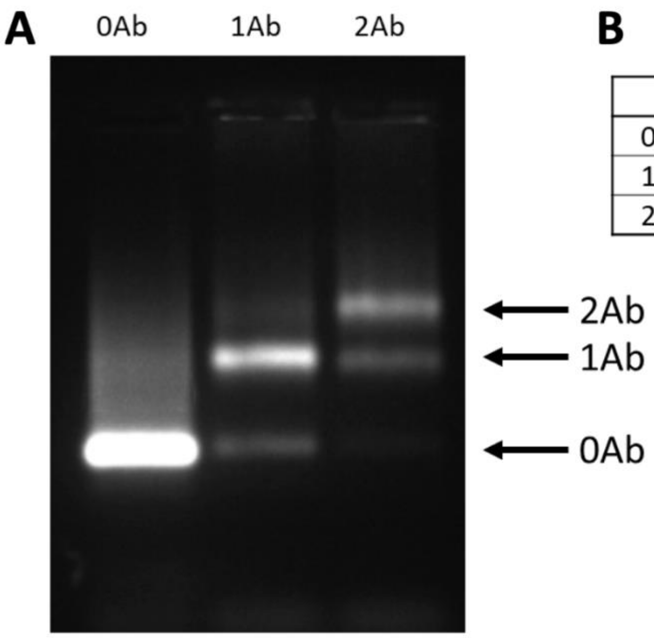


**B** Yield [%]

| | 1 Overhang | 2 Overhangs |
|---|---|---|
| 0Ab | 0.19 | 0.04 |
| 1Ab | 0.79 | 0.25 |
| 2Ab | | 0.70 |

**Figure S3:** Gel analysis of antibody labeling of DNA origami. A) Agarose gel electrophoresis analysis of labeling 8HB DNA origami with 1 or 2 antibodies. B) Intensity analysis of the gel bands indicates labeling efficiencies of 70% or better.

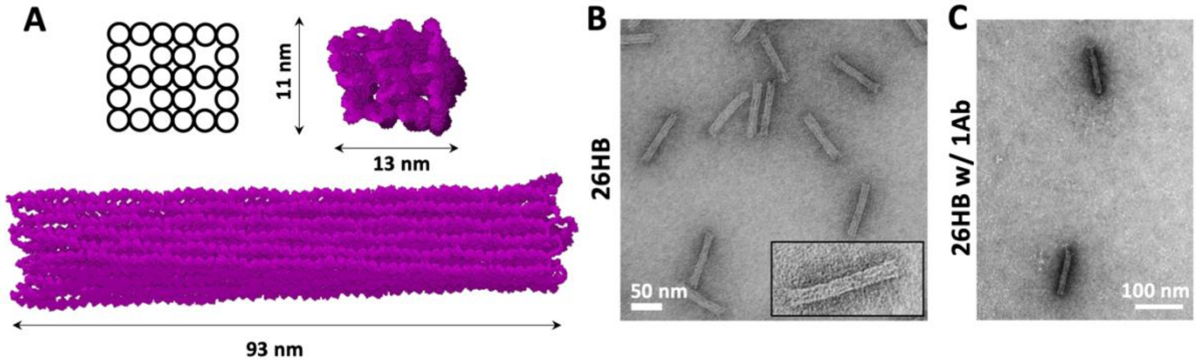


**Figure S4:** Simulation and characterization of 26HB. A) Simulated 26HB design showing side view (bottom) and front view (top right). The cross-section is shown at top left with circles representing dsDNA helices. B) TEM image of 26HB. C) TEM image of 26HB with 1 anti Pol II antibody.

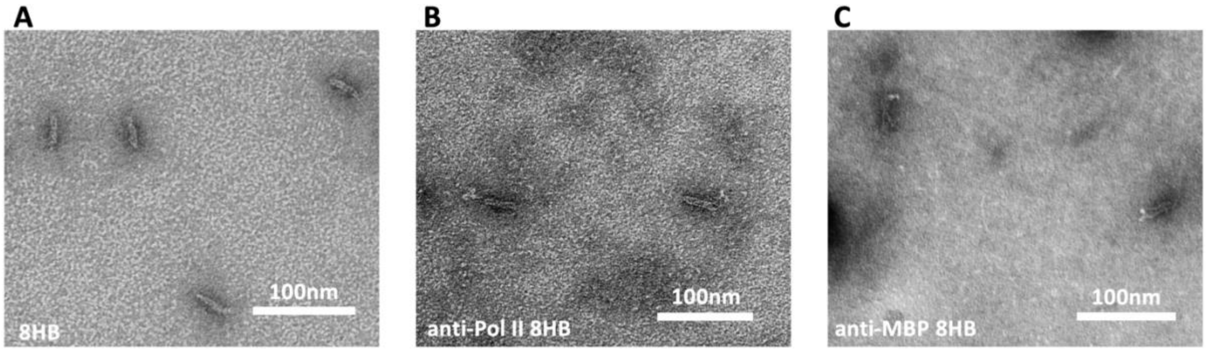


**Figure S5:** Additional TEM characterization of 8HB. A) TEM image of 8HB structures alone. B) TEM image of 8HB with 1 anti-Pol II antibody attached (anti Pol II 8HB). C) TEM image of 8HB with 1 anti-MBP antibody attached (anti MBP 8HB).

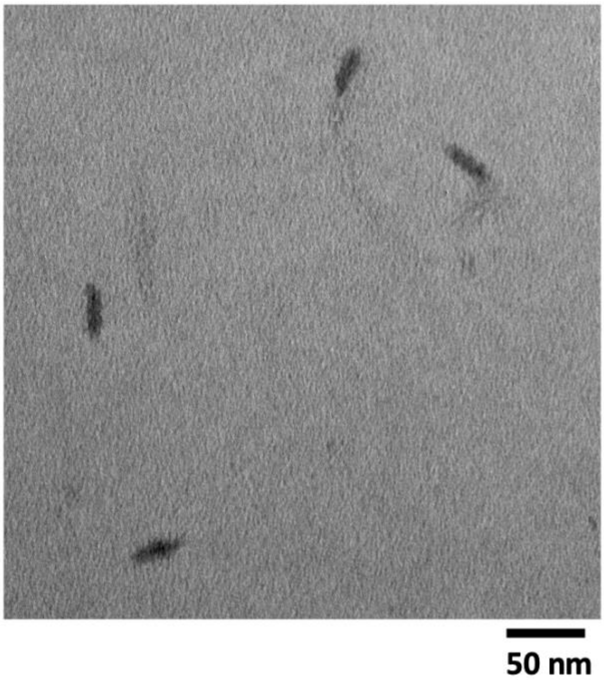


**Figure S6:** TEM image of 8HB after incubation in cell culture media at 37 $^{o}$C for 24 hr illustrating intact 8HB structures.

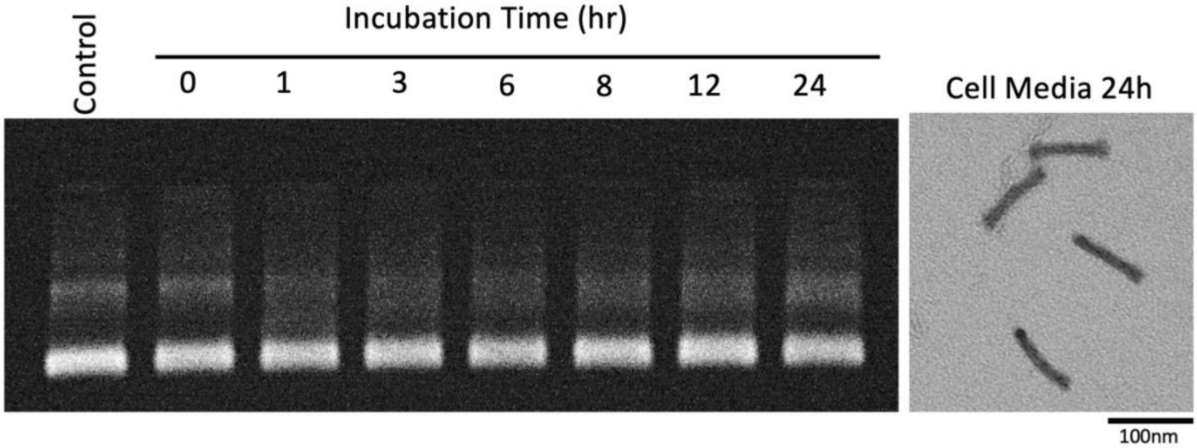


**Figure S7:** Stability of 26HB-8Cy5 in cell culture media. The left image shows agarose gel electrophoresis (Cy5 fluorescence emission) analysis of 26HB-8Cy5 structures incubated in cell culture media at 37 °C for varying time points up to 24hr showing the 26HB-8Cy5 remains stable. TEM imaging (right) of the 24 hr sample confirms intact structures.

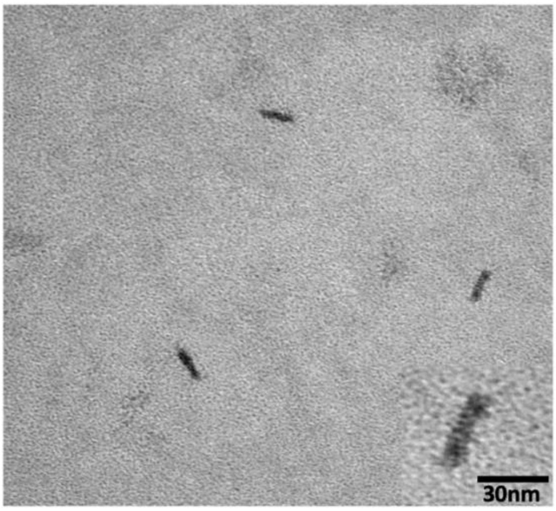


**Figure S8:** Additional TEM characterization of 8HB-8Cy5 incubated for 24 hr in U2OS nuclear extract at 37 °C.

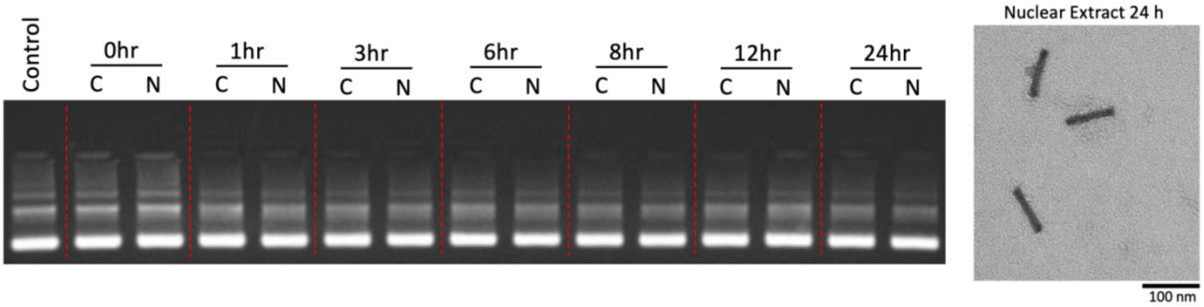


**Figure S9:** Stability of 26HB-8Cy5 in U2OS celll extracts. The left image shows agarose gel electrophoresis (Cy5 fluorescence emission) analysis of 26HB-8Cy5 structures incubated at 37 °C in U2OS cytoplasmic (C) or nuclear (N) extracts for varying time points up to 24hr showing the 26HB-8Cy5 remains stable. TEM imaging (right) of the sample incubated in nuclear extract for 24 hr at 37 °C confirms intact structures.

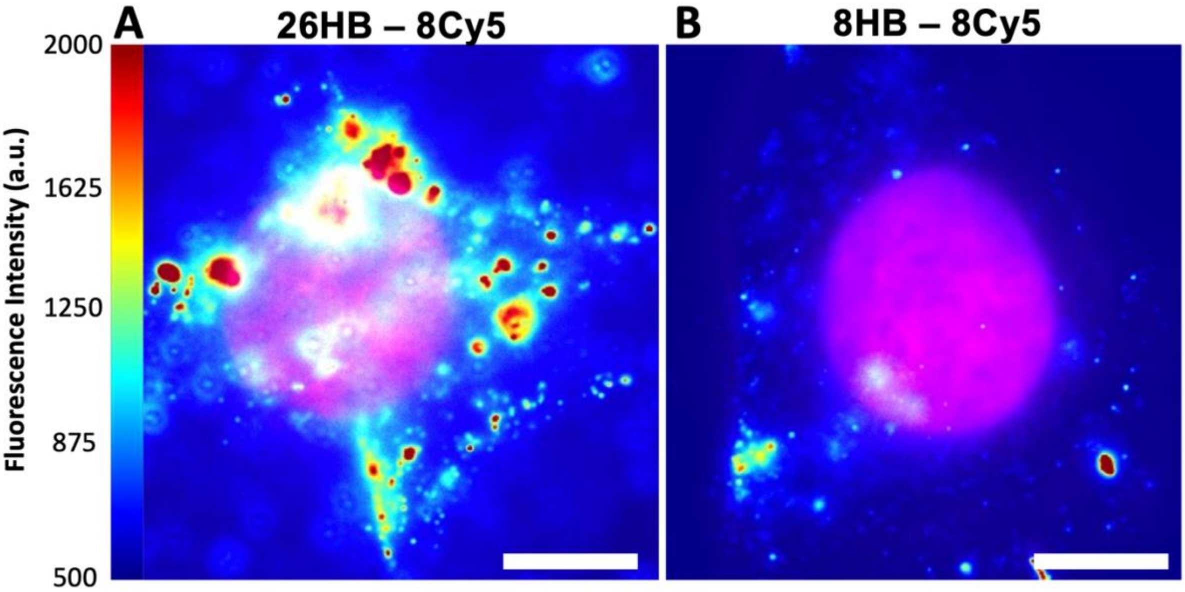


**Figure S10:** HILO imaging of electroporated 26HB-8Cy5 and 8HB-8Cy5 reveals cytosolic aggregation of the 26HB. A) Fluorescence imaging revealed significant aggregation of the 26HB-8Cy5 structure electroporated into the cytoplasm of U2OS cells. Very few smaller spots indicative of individual origami were observed. B) Fluorescence imaging of the 8HB-8Cy5 sample electroporated into the cytoplasm of U2OS cells reveal mostly smaller spots indicative of well dispersed DNA origami structures. Nuclear signal (magenta) from Hoechst stain imaged in separate channel. Heat bar = fluorescence intensity, Scale bars = 10 μm. Since the 26HB exhibited significant aggregation we focused the remainder of our experiments on the 8HB structures.

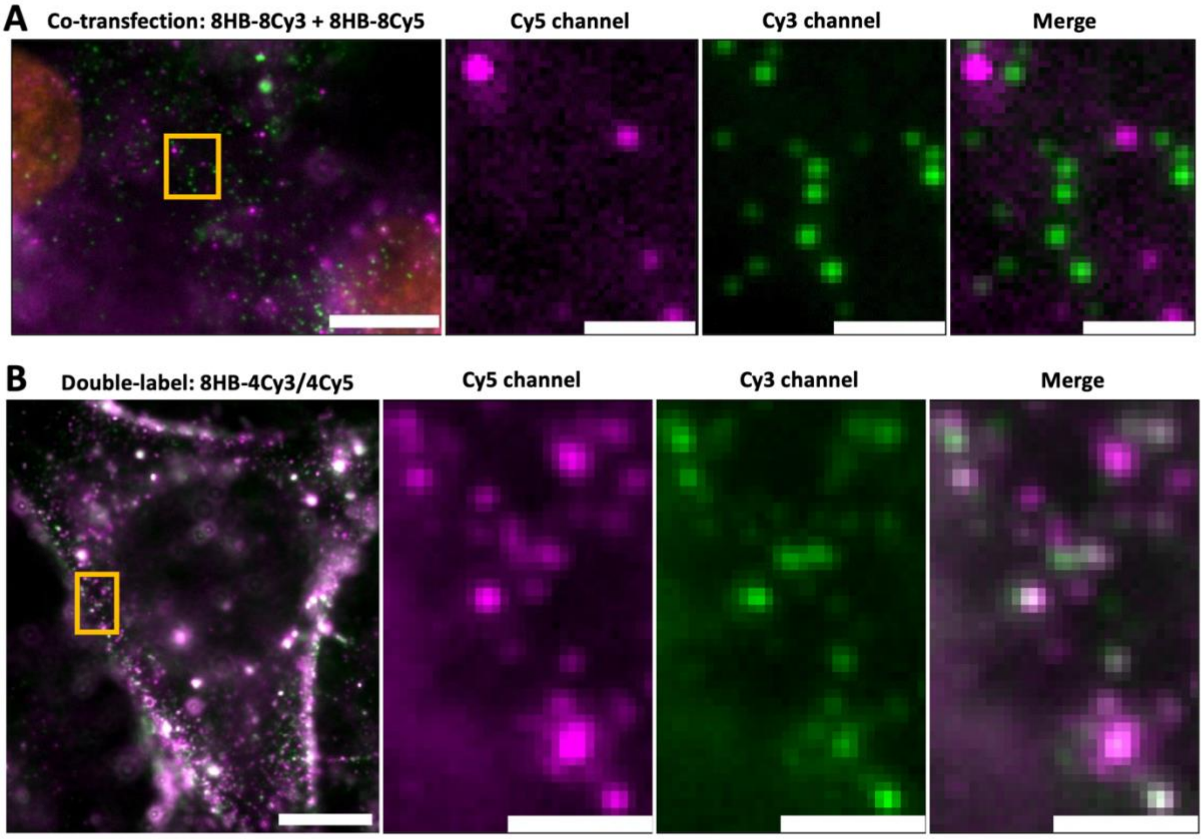


**Figure S11:** Stability of fluorophore-labeled DO after transfection into U2OS cells. A) HILO image of U2OS cells transfected with a combination of 8HB-8Cy5 and 8HB-8Cy3. Nuclear signal is Hoechst stain imaged in separate channel from Cy3 and Cy5 signals. The images reveal no significant observed aggregation. B) HILO image of U2OS cell transfected with DNA origami nanostructure labeled with both Cy5 and Cy3 fluorophores (8HB-4Cy3/4Cy5). The images show most spots exhibit both Cy3 (green) and Cy5 (magenta) fluorescence. The co-localization was quantified in terms of the average Pearson correlation coefficient. The 8HB structures exhibited correlation coefficients of $r = 0.82 \pm 0.10$ for the double-labeled 8HB (8HB-4Cy5/4Cy3) and $r = 0.34 \pm 0.14$ for the co-transfected single labeled 8HB (8HB-8Cy5 plus 8HB-8Cy3). Large field of view scale bars = 10μm, Zoom scale bars = 2μm.

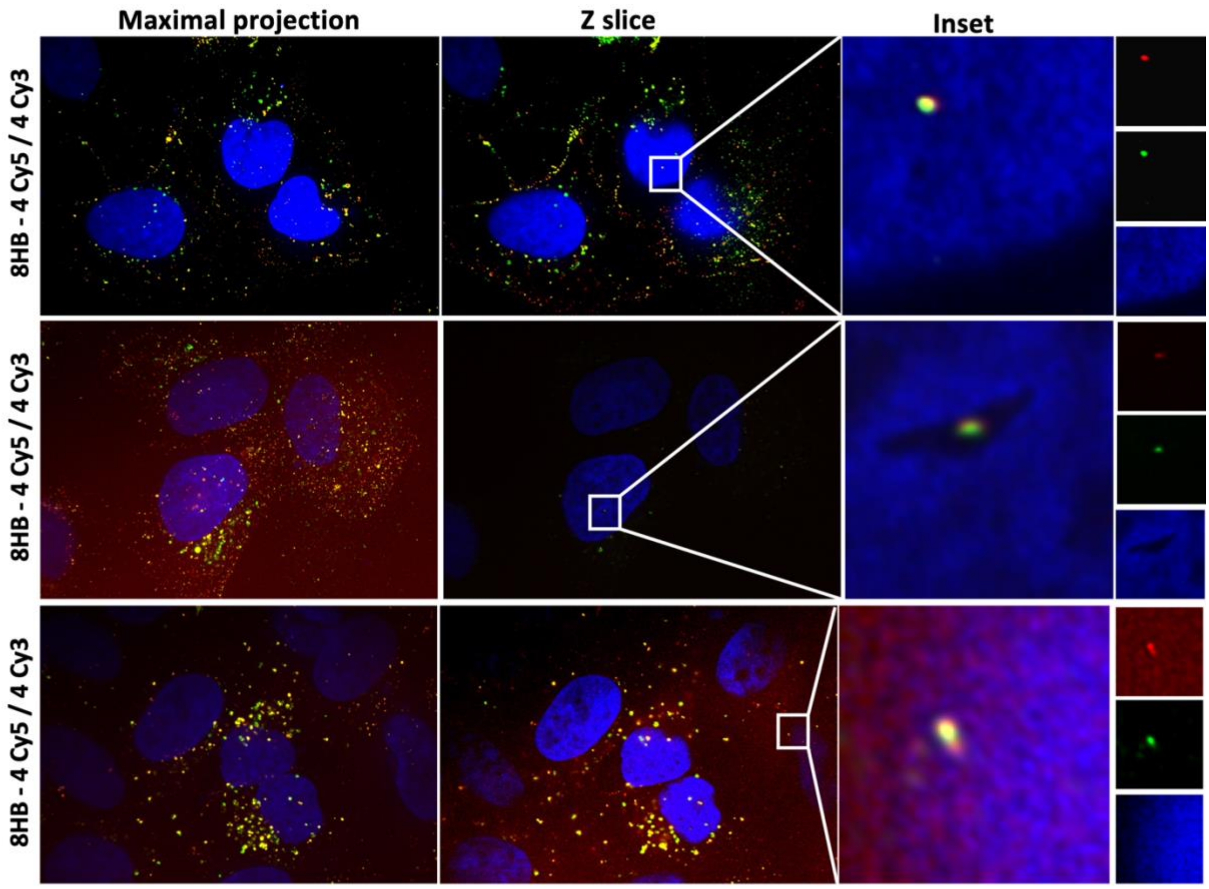


**Figure S12:** Additional iSIM images of 8HB-4Cy5/4Cy3 electroporated into U2OS cells. The left column shows maximal intensity projections and the middle column shows a single z-slize at the plane of largest nuclear area. The right column shows zoomed in views highlighting examples of co-localization of Cy3 and Cy5 signals from 8HB structures inside the nuclear. The right side shows the three separate channels of the zoom in images.

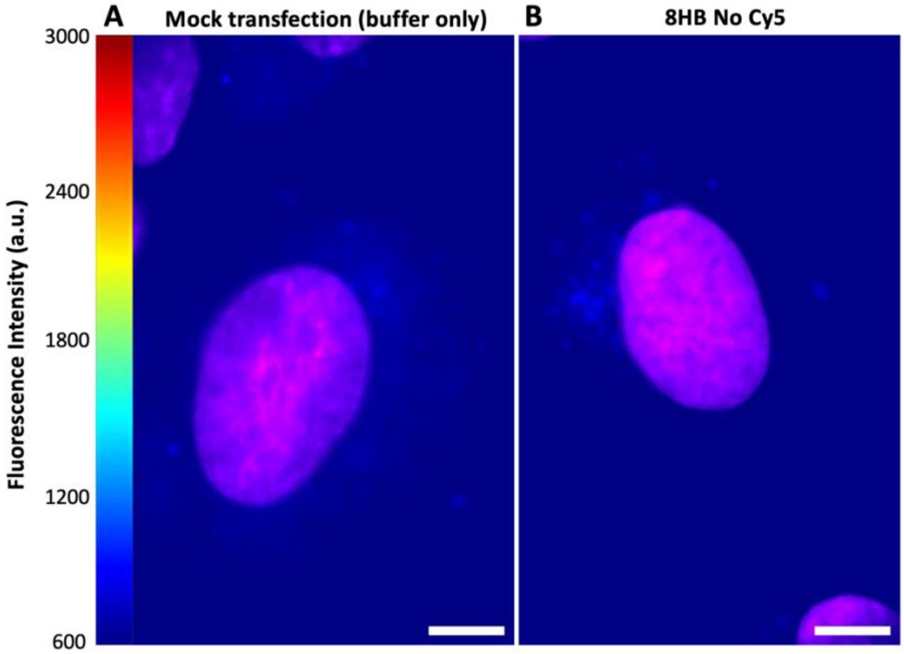


**Figure S13:** Transfection controls. A) HILO image of U2OS cell transfected with electroporation buffer only. B) HILO image of U2OS cell transfected with 8HB structures containing no fluorescent label. Nuclear signal (magenta) from Hoechst stain imaged in separate channel. Heat bar = fluorescence intensity of origami channel, Scale bars = 10μm.

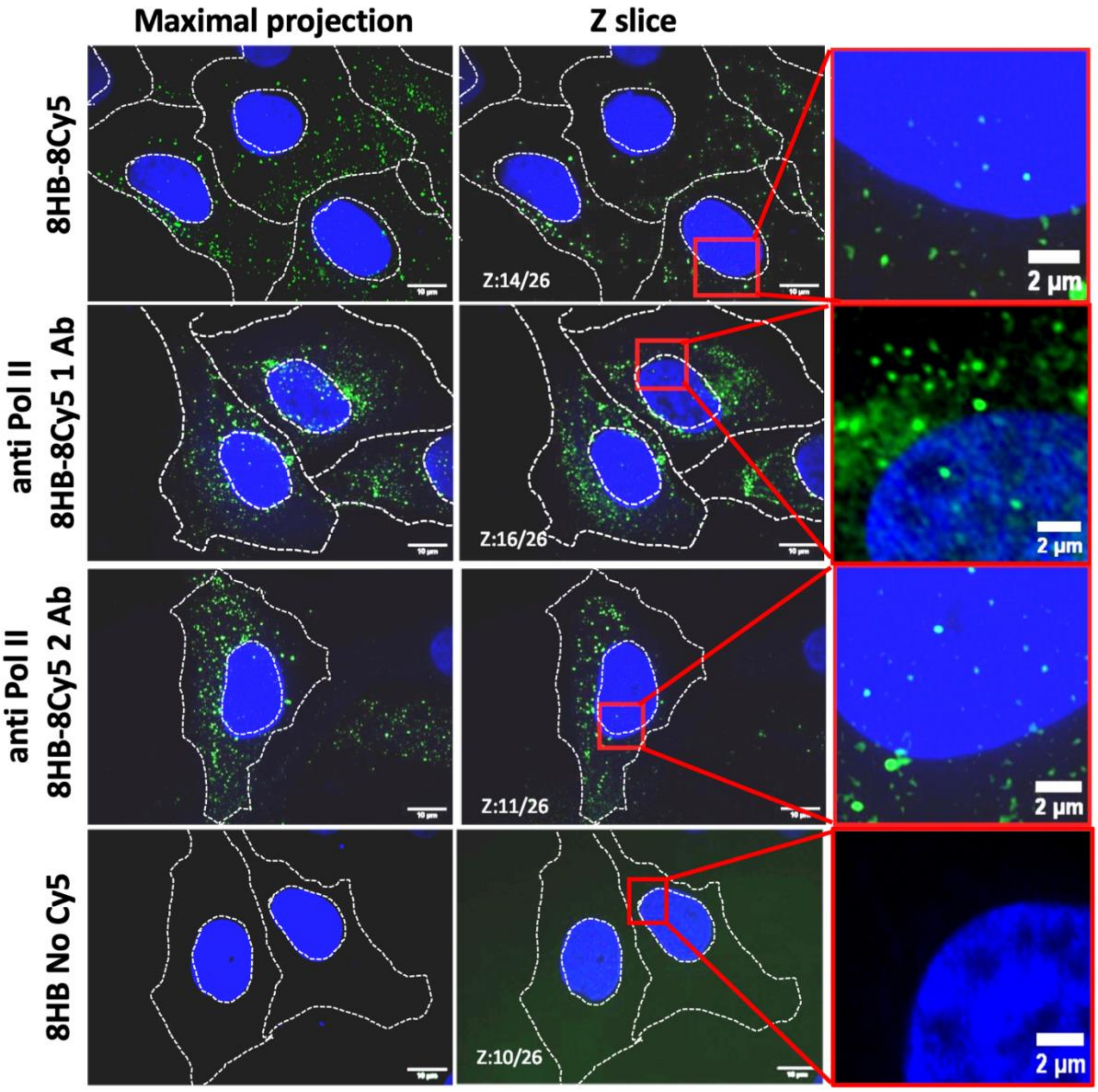


**Figure S14:** Confocal microscopy analysis of 8HB DO (green) localization in U2OS cells. Image acquisition was set at the middle of the sample based on DAPI / Hoeschst staining (blue) and Z-stacks of 5 µm of total thickness were acquired with 0.2 um step size. Images were deconvoluted with microvolution plugin in ImageJ software. Maximal projection of all images in the stack. The top row shows the 8HB-8Cy5 (no antibody labeling). The second row shows the 8HB-8Cy5 labeled with one anti-Pol II antibody (anti Pol II 8HB-8Cy5 1 Ab). The third row shows the 8HB-8Cy5 labeled with two anti-Pol II antibodies (anti Pol II 8HB-8Cy5 2 Ab). The bottom row shows the 8HB structure with no antibody or fluorophore labeling. Large field of view scale bars are 10 µm. The left column shows maximal intensity projections, the middle column shows a single z-slice at the largest nuclear area. The right column shows a zoom in of the single z slice highlighting examples of 8HB in the nucleus.

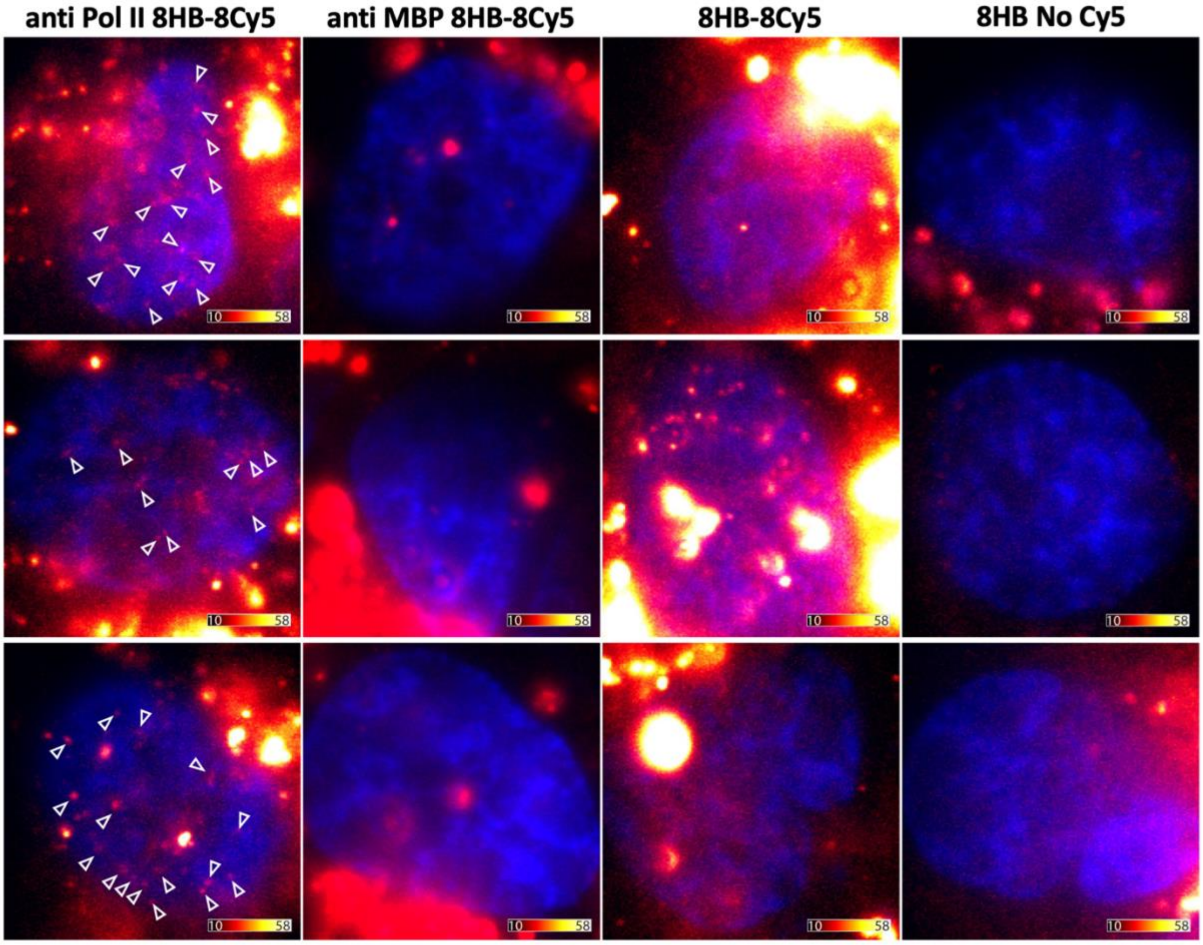


**Figure S15:** Nuclear delivery of DNA origami in live cells. Representative images of live U2OS cells 24 hr after electroporation with non-functionalized 8HB structures (right) or 8HB functionalized with either one anti-Pol II antibody and labelled with Cy5 (anti-Pol II-8HB-8Cy5; left), or 8HB functionalized with one anti-MBP antibody and Cy5 labelled (anti-MBP-8HB-8Cy5; 2nd from left), or 8HB labelled with Cy5 (8HB-8Cy5; 2nd from right). White arrowheads point to nuclear particles representing single origami structures. Nuclear Hoechst33342 staining is shown in blue. Color bars are 5 µm long and indicate the fluorescence intensity range of the Cy5 signal (on a scale of x1000).

**Supplementary video 1:** Nuclear delivery of DNA origami in live cells. U2OS cells were electroporated with non-functionalized 8HB origami structures (8HB) or 8HB functionalized with either 1 anti-Pol II antibody and 8 Cy5 molecules (left), or 1 anti-MBP antibody and 8 Cy5 molecules (2nd from left), or 8 Cy5 molecules (2nd from right). Cells were stained with Hoechst and imaged using live HILO microscopy. Time laps of the first 100 frames are shown. White lines denote the outlines of nuclei. Scale bars are 5 µm.

**Supplementary video 2:** Nuclear delivery of different antibody-conjugated 8HB origami structures in live U2OS cells. Representative examples of U2OS cells that were electroporated with 8HB origami structures functionalized with either 1 anti-Pol II antibody and 8 Cy5 molecules (top row) or with 1 anti-MBP antibody and 8 Cy5 molecules (bottom row). Cells were stained with Hoechst and imaged using live HILO microscopy. Time lapse of the firs 100 frames are shown. White lines denote the outlines of nuclei. Scale bars are 5 µm.
